# Single cell ‘omic profiles of human aortic endothelial cells *in vitro* and human atherosclerotic lesions *ex vivo* reveals heterogeneity of endothelial subtype and response to activating perturbations

**DOI:** 10.1101/2023.04.03.535495

**Authors:** Maria L. Adelus, Jiacheng Ding, Binh T. Tran, Austin C. Conklin, Anna K. Golebiewski, Lindsey K. Stolze, Michael B. Whalen, Darren A. Cusanovich, Casey E. Romanoski

## Abstract

**Objective:** Endothelial cells (ECs), macrophages, and vascular smooth muscle cells (VSMCs) are major cell types in atherosclerosis progression, and heterogeneity in EC sub-phenotypes are becoming increasingly appreciated. Still, studies quantifying EC heterogeneity across whole transcriptomes and epigenomes in both *in vitro* and *in vivo* models are lacking.

**Approach and Results:** To create an *in vitro* dataset to study human EC heterogeneity, multiomic profiling concurrently measuring transcriptomes and accessible chromatin in the same single cells was performed on six distinct primary cultures of human aortic ECs (HAECs). To model pro-inflammatory and activating environments characteristic of the atherosclerotic microenvironment *in vitro*, HAECs from at least three donors were exposed to three distinct perturbations with their respective controls: transforming growth factor beta-2 (TGFB2), interleukin-1 beta (IL1B), and siRNA-mediated knock-down of the endothelial transcription factor ERG (siERG). To form a comprehensive *in vivo/ex vivo* dataset of human atherosclerotic cell types, meta-analysis of single cell transcriptomes across 17 human arterial specimens was performed. Two computational approaches quantitatively evaluated the similarity in molecular profiles between heterogeneous *in vitro* and *in vivo* cell profiles. HAEC cultures were reproducibly populated by 4 major clusters with distinct pathway enrichment profiles: EC1-angiogenic, EC2-proliferative, EC3-activated/mesenchymal-like, and EC4-mesenchymal. Exposure to siERG, IL1B or TGFB2 elicited mostly distinct transcriptional and accessible chromatin responses. EC1 and EC2, the most canonically ‘healthy’ EC populations, were affected predominantly by siERG; the activated cluster EC3 was most responsive to IL1B; and the mesenchymal population EC4 was most affected by TGFB2. Quantitative comparisons between *in vitro* and *in vivo* transcriptomes confirmed EC1 and EC2 as most canonically EC-like, and EC4 as most mesenchymal with minimal effects elicited by siERG and IL1B. Lastly, accessible chromatin regions unique to EC2 and EC4 were most enriched for coronary artery disease (CAD)-associated SNPs from GWAS, suggesting these cell phenotypes harbor CAD-modulating mechanisms.

**Conclusion:** Primary EC cultures contain markedly heterogeneous cell subtypes defined by their molecular profiles. Surprisingly, the perturbations used here, which have been reported by others to be involved in the pathogenesis of atherosclerosis as well as induce endothelial-to-mesenchymal transition (EndMT), only modestly shifted cells between subpopulations, suggesting relatively stable molecular phenotypes in culture. Identifying consistently heterogeneous EC subpopulations between *in vitro* and *in vivo* models should pave the way for improving *in vitro* systems while enabling the mechanisms governing heterogeneous cell state decisions.

## INTRODUCTION

Endothelial Cells (ECs) in the vascular endothelium maintain hemostasis, mediate vasodilation, and regulate the migration of leukocytes into tissues during inflammation. Dysfunctions of the endothelium are a hallmark of the aging process and are also an important feature of diseases including atherosclerosis. Atherosclerosis is an inflammatory process fueled by cholesterol and leukocyte accumulation in the sub-endothelial layer of arteries. It is the underlying pathobiology of ischemic heart disease and the leading cause of morbidity and mortality worldwide due to heart attack and stroke (1–3). Atherosclerosis of the coronary arteries is estimated to be about 50% genetic with hundreds of genomic loci contributing to genetic risk (4–6). A major opportunity for better understanding the molecular basis for how disease progresses lie in identifying the genomic and downstream functions impaired by risk variants in disease-relevant cell types. Genetic studies are increasingly suggesting that a significant proportion of genetic risk for atherosclerosis is encoded in perturbed functions of vascular ECs (5–7).

Single cell sequencing technologies have begun to characterize the extent of EC molecular diversity *in vitro* and *in vivo* (8–19). Genetically engineered, lineage traced mouse models have also been instrumental for identifying which cells in atherosclerotic plaques arose from EC origin. These studies have demonstrated that many cells of EC origin in plaques lack canonical EC marker genes and luminal location (20, 21). As many as one-third of mesenchymal-like cells in plaques have been reported to be of endothelial origin (20) suggesting that phenotypic transition from endothelial to mesenchymal (EndMT) is a feature of atherosclerosis; however, whether EndMT is a cause or bystander of atherogenesis or plaque rupture is not fully understood. Although lineage tracing is not possible in humans, immunocytochemical techniques suggest that EC heterogeneity is prevalent in atherosclerotic vessels. These studies have described an unexpectedly large number of cells co-expressing pairs of endothelial and mesenchymal proteins, including fibroblast activating protein/von Willebrand factor (FAP/VWF), fibroblast-specific protein-1/VWF (FSP-1/VWF), FAP/platelet-endothelial cell adhesion molecule-1 (CD31 or PECAM-1), FSP-1/CD31 (20), phosphorylation of TGFB signaling intermediary SMAD2/FGF receptor 1 (p-SMAD2/FGFR1) (22), and α-smooth muscle actin (αSMA)/PECAM-1 (23). An important implication of this result is that the use of canonical EC markers to isolate or identify ECs will likely omit certain EC populations. The extent of EC molecular and functional heterogeneity within a tissue during homeostasis and during disease is not well understood. One notable study exemplifying EC heterogeneity demonstrated that the EC-marker gene von Willebrand Factor (*VWF*) was expressed only in a subset of ECs from the same murine vessel, and the penetrance of *VWF* expression across ECs was tissue-specific (24). In a related study, expression of the leukocyte adhesion molecule *VCAM-1* was found to be upregulated by the pro-inflammatory cytokine tumor necrosis factor alpha (TNFa) only in some of the ECs of a monolayer (25). In both studies, variability in DNA methylation on CpG dinucleotides at the gene promoters negatively correlated with *VWF* and *VCAM-1* expression. These findings raise the question as to how many molecular programs exist within ECs of a same tissue or culture, how this heterogeneity influences response to cellular perturbations, and what factors regulate these cellular states.

There are notable benefits and limitations for studying heterogeneity using *in vitro* and *in vivo* approaches in atherosclerosis research. *In vitro* approaches provide unique opportunities for interrogating consequences of genetic and chemical perturbations in highly controlled environments and are adept at identifying mechanistic relationships on accelerated timelines. *In vivo* approaches benefit from the complexity of the crosstalk among all cell types and tissues of the organism and are adept for identifying how perturbations manifest in living systems. It reasons that the integration of results from both approaches will best accelerate discovery. However, comprehensive analysis comparing heterogeneity of vascular ECs observed *in vivo* and *in vitro* remains unexplored. In the current study we performed meta-analysis on four human *in/ex vivo* single cell transcriptomic datasets (26–29), containing 17 arterial samples, from mild-to-moderate calcified atherosclerotic plaques to evaluate the ability of the *in vitro* EC models to recapitulate molecular signatures observed in human atherosclerosis.

Human aortic endothelial cells (HAECs) are among the most appropriate cell type for *in vitro* modeling of the arterial endothelium in atherosclerosis research insofar as they are human cells, they are more readily available than coronary artery ECs, they are not of venous origin like human umbilical vein ECs, and they can be isolated from explants of healthy donor hearts during transplantation. We set forth in the current study to quantify heterogeneity among HAECs using multimodal sequencing that simultaneously measures transcripts using RNA-seq and accessible chromatin using ATAC-seq from the same barcoded nuclei. To provide estimates for heterogeneity due to genetic background, we molecularly phenotyped HAECs from six genetically distinct human donors. We also quantified single cell responses to three perturbations known to be important in EC biology and atherosclerosis. The first was activation of transforming growth factor beta (TGFB) signaling, which is a hallmark of phenotypic transition and a regulator of EC heterogeneity (20, 30). The second was stimulation with the pro-inflammatory cytokine interleukin-1 beta (IL1B), which has been shown to model inflammation and EndMT *in vitro* (31–35), and whose inhibition reduced adverse cardiovascular events in a large clinical trial (36). The third perturbation utilized in our study was knock-down of the ETS related gene (*ERG*), which encodes a transcription factor of critical importance for EC fate specification and homeostasis (37–41).

Lastly, we examine whether epigenetic landscapes among heterogeneous EC subtypes observed in this study were differentially enriched for coronary artery disease (CAD) genetic risk variants. Taken together, this study provides evidence that EC heterogeneity is prevalent *in vivo* and *in vitro* and that not all ECs respond similarly to activating perturbations.

## RESULTS

### EC Single Cell Transcriptomic Profiles Reveal a Heterogeneous Population

To systematically uncover the heterogeneity of molecular landscapes in ECs at single cell resolution, we cultured primary HAECs isolated from luminal digests of ascending aortas from six de-identified heart transplant donors at low passage (passage 3-6) (42) (**Figure 1A**). Using the 10X Genomics multiome kit (43), single nucleus mRNA expression (snRNA-seq) and chromatin accessibility (snATAC-seq) data were collected simultaneously for a total of 15,220 nuclei after stringent quality control (**Methods**). RNA and ATAC data were integrated separately by treatment condition and then with each other as reported previously (**Methods**) (44). snRNA-seq libraries were sequenced to a median depth of 29,732-84,476 reads and 2,481-3,938 transcripts per nucleus (**Table S1** and **Table S2** in the **Data Supplement**). Five distinct EC subtypes (EC1, EC2, EC3, EC4, and EC5) were detected from the fully integrated dataset, which included all donors, treatments, and data types (**Figure 1B**). Subtypes EC1 and EC3 comprised cells from all donors, whereas EC2 and EC4 contained cells from most donors, and EC5 was nearly exclusively populated by cells from a single donor (**Figure 1C**; **Table S3** in the **Data Supplement**). Because we do not observe EC5 across multiple individuals, we chose not to focus additional analysis on this subtype. Pathway enrichment of marker genes revealed EC1 to exhibit an angiogenic phenotype (WP4331, p-value 4.0×10^-9^; GO:0038084, p-value 1.5×10^-9^) with enriched transcripts including *KDR*, *GAB1*, *PGF*, and *NRP2* (**Figure 1D-G**, **Figure S1A** in the **Data Supplement**). EC2 was enriched in proliferation (GO:1903047, p-value 7.4×10^-35^) with characteristic markers *CENPE*, *CENPF*, *KIF11*, *KIF4A* and *TOP2A* (**Figure 1D-G**, **Figure S1A** in the **Data Supplement**). EC3 displayed enrichment in “regulation of smooth muscle cell proliferation” (GO:0048660; p-value 1.1×10^-10^) (**Figure 1F**). From the top 200 differentially expressed genes (DEGs) for EC3 we observed additional pathways enriched, including NABA CORE MATRISOME (M5884; p-value 1×10^-34^) and locomotion (GO:0040011; p-value 1.2×10^-15^), suggesting an activated mesenchymal-like phenotype (**Figure S1B-C** in the **Data Supplement**). A fourth subset, EC4, demonstrates enrichment in ECM organization (GO:0097435; p-value 3.2×10^-19^), a process characteristic of mesenchymal cells, with distinctive expression of collagen genes, including *COL1A1*, *COL1A2*, *COL3A1*, and *COL5A1* (**Figure 1D-G**, **Figure S1A** in the **Data Supplement**) (45, 46). Top marker genes and pathways for each EC subtype are in **Table S4**-**5** in the **Data Supplement**. These observations are in line with previous reports of angiogenic, proliferative, mesenchymal, and pro-coagulatory EC subtypes within *ex vivo* models (9, 10, 14, 19, 47) and underscore the heterogeneity of transcriptomic profiles in cultured HAECs.

**Figure 1.**
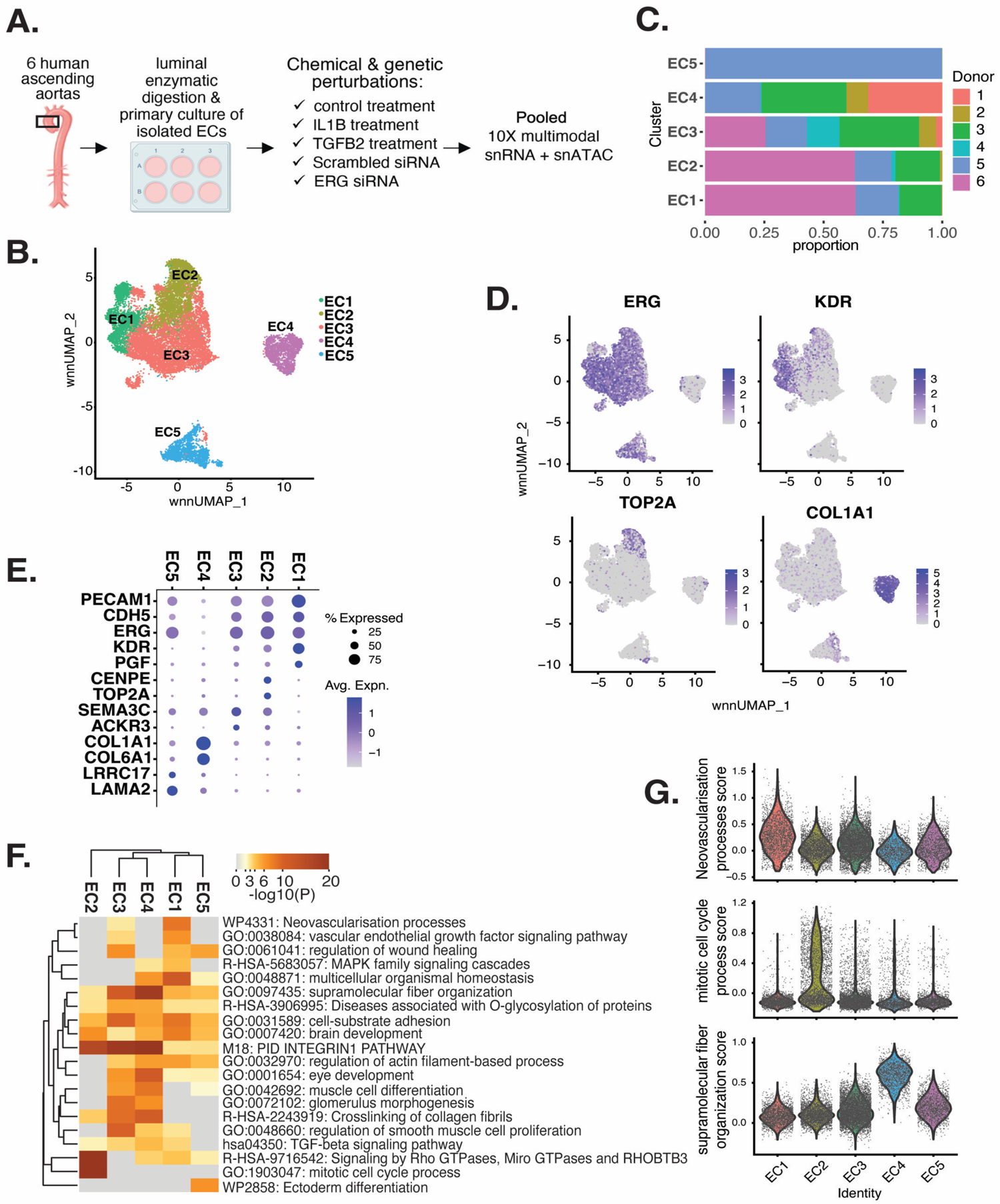
HAEC transcriptomic profiling discover a heterogenous cell population. **(A)**, Schematic diagram of the experimental design. ECs were isolated from six human heart transplant donor’s ascending aortic trimmings and treated with IL1B, TGFB2, or siERG (ERG siRNA) for 7 days **(B)**, Weighted Nearest Neighbor UMAP (_WNN_UMAP) of aggregate cells from all perturbations and donors is shown. Each dot represents a cell, and the proximity between each cell approximates their similarity of both transcriptional and epigenetic profiles. Colors denote cluster membership. **(C)**, The proportion of cells from each donor for each EC subtype. **(D)**, Gene expression across top markers for each cluster including pan EC (*ERG*), EC1 (*KDR*), EC2 (*TOP2A*), and EC4 (*COL1A1*). **(E)**, Top markers for pan EC (*PECAM1*, *CDH5*, *ERG*), EC1 (*KDR*, *PGF*), EC2 (*CENPE*, *TOP2A*), EC3 (*SEMA3C*, *ACKR3*), EC4 (*COL1A1*, *COL6A1*), and EC5 (*LRRC17*, *LAMA2*). The size of the dot represents the percentage of cells within each EC subtype that express the given gene, while the shade of the dot represents the level of average expression (“Avg. Expn.” in the legend). **(F)**, Heatmap of pathway enrichment analysis (PEA) results from submitting top 200 differentially expressed genes (DEGs; by ascending p-value) between EC subtypes. Rows (pathways) and columns (EC subtypes) are clustered based on -Log_10_(P) **(G)**, Violin plots of top Metascape pathway module scores across EC subtypes. Module scores are generated for each cell barcode with the Seurat function AddModuleScore().

### EC Subtypes Exhibit Distinct Open Chromatin Profiles and Enriched Motifs

To investigate how different transcriptional signatures across ECs correspond to distinct chromatin states, we utilized the snATAC-seq portion of the multiome dataset. The snATAC-seq data were sequenced to a median depth of 22,939-126,122 reads with 3,480-19,259 peaks called per nucleus (**Table S2** and **S6** in the **Data Supplement**). Of 204,904 total identified peaks, 13,731 were differential across subtypes, with 79 to 8,091 peaks uniquely accessible per EC subtype (**Table S8** in the **Data Supplement**). Over 80% of total peaks were intergenic or intronic (Figure 2A-B) and most unique peaks were from EC2 and EC4.

**Figure 2.**
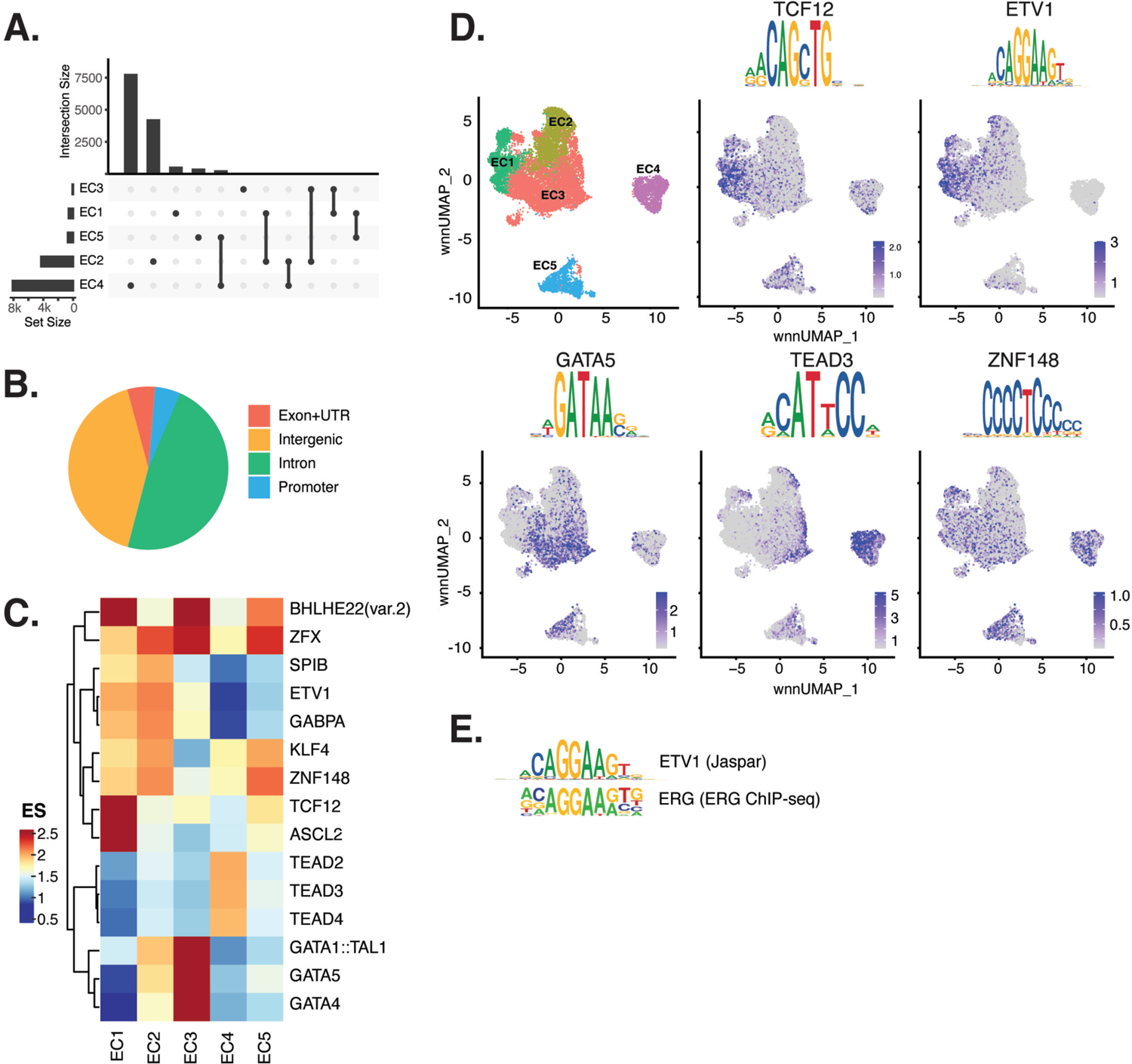
ECs have epigenetically distinct cell states. **(A)**, Upset plot of differential peaks across EC subtypes. Intersection size represents the number of genes at each intersection, while set size represents the number of genes for each EC subtype. **(B)**, Genomic annotation for the complete peak set. **(C)**, Heatmap of top transcription factors (TFs) from motif enrichment analysis for marker peaks in each EC subtype. Top TFs for each EC subtype are selected based on ascending p-value. Rows (TFs) and columns (EC subtype) are clustered based on enrichment score (ES). **(D)**, Feature plots and position weight matrices (PWMs) for top TF binding motifs for EC1 (TCF12), EC2 (ETV1), EC3 (GATA5), and EC4 (TEAD3). Per-cell motif activity scores are computed with chromVAR, and motif activities per cell are visualized using the Signac function FeaturePlot. **(E)**, PWMs comparing Jaspar 2020 ETV1 motif to ERG motif reported in Hogan et al.

Transcription factor (TF) motif enrichment analysis using Signac (48) was performed on Differentially Accessible Regions (DARs) per EC subtype (Figure 2C). It is important to note that TFs within a TF family may share DNA-binding motifs and may not be distinguished by motifs alone. As a result, TF names from the Jaspar database (49) indicate the TF family. We find the basic helix-loop-helix (*bHLH*) motif defined by the core sequence CANNTG enriched in EC1 peaks, including enrichments for ASCL2 (adjusted p-value 3.9×10^-50^), TCF12 (adjusted p-value 1.7×10^-21^), and BHLHE22(var.2) (adjusted p-value 5.7×10^-48^) (Figure 2C-D). ETS motifs, including ETV1 (adjusted p-value 3.2×10^-42^ and 5.3×10^-249^, for EC1-2, respectively), SPIB (adjusted p-value 7.9×10^-22^ and 2.5×10^-236^, respectively), and GABPA (adjusted p-value 2.7×10^-41^ and 4.3×10^-244^, respectively), were also enriched in EC1 as well as in EC2 peaks. These data are consistent with known roles for ETS TFs, including ERG and FLI1, in governing angiogenic and homeostatic endothelial phenotypes (50). Given that *ERG* expression (Figure 1E) correlated with incidence of the ETS motif in open chromatin (Figure 2D) across the nuclei, ERG is likely driving the EC1-2 sub-phenotypes. The near-exact match in motifs between the ETV1 motif position weight matrix in Jaspar and the *de novo* enriched motif from ERG ChIP-seq in human aortic ECs (41) further supports this conclusion (Figure 2E). In addition to ETS motifs, EC2 was enriched in ZFX (adjusted p-value 4.2×10^-86^) and ZNF148 (adjusted p-value 1.1×10^-126^), which are C2H2 zinc finger motifs. C2H2 zinc finger motifs, as well as KLF4 (adjusted p-value 5.4×10^-32^ and 8.4×10^-135^, for EC1-2, respectively), also show enrichment in EC1 and EC2. EC3 peaks are enriched for GATA motifs including GATA4 (adjusted p-value 3.1×10^-8^), GATA5 (adjusted p-value 8×10^-11^), GATA1::TAL1 (adjusted p-value 1.8×10^-6^), and bHLH motif BHLHE22(var.2) (adjusted p-value 0.01). EC4 open regions were uniquely enriched for TEA domain (TEAD) factors comprised of motifs named TEAD2 (adjusted p-value 1.2×10^-238^), TEAD3 (adjusted p-value 2.1×10^-306^), and TEAD4 (adjusted p-value 6.9×10^-252^) (Figure 2C-D). Notably, TEAD factors have been found as enriched in vascular smooth muscle cells (VSMCs) (29, 51), which is consistent with EC4 having the most mesenchymal phenotype of our EC subtypes.

Taken together, these data demonstrate that EC1 and EC2 are the subtypes most canonically like ‘healthy’ or angiogenic ECs insofar as they exhibit ETS motif enrichments. Additionally, we conclude that EC4 is the most mesenchymal EC insofar as it exhibits TEAD factor enrichments.

### EC activating perturbations modestly shift cells into the EC3 subtype

Embedded in the dataset of this study were three experimental conditions known to promote EndMT along with their respective controls. Each experimental condition was administered to between three and five genetically distinct HAEC cultures. The conditions included 7-day exposure to IL1B (10 ng/ml), 7-day exposure to TGFB2 (10 ng/mL), and 7-day siRNA-mediated knock-down of ERG (siERG). The control for IL1B and TGFB2 treatments was 7-day growth in matched media lacking cytokine and the control for the siERG condition was transfection with scrambled RNA.

The UMAP presented in Figure 1 includes all the nuclei profiled across donors and conditions. We hypothesized that EC4, the most mesenchymal cluster, would be enriched for cells exposed to IL1B, TGFB2, and/or siERG relative to the controls thereby consistent with the hypothesis that the EC4 subtype were a consequence of EndMT. Detailed in Figure 3A-B are the relative proportions of cells from each experimental condition and donor by cluster. Contrary to our hypothesis, the EC4 cluster was not enriched for cells that were treated with cytokine or siERG relative to the controls; in fact, there is a non-statistically significant trend for decreased numbers of EC4 cells from these conditions relative to controls insofar as all the donors with cells in EC4 show diminished proportions upon perturbation (Figure 3). The one cluster exhibiting increased proportions of cells upon perturbations was EC3, with 3 of 4 EC IL1B-exposed donors having increased proportions in EC3 (p = 0.08 by 2-sided paired t-test; Figure 3A), 4 of 5 TGFB2-exposed donors having increased proportions (p = 0.04 by 2-sided paired t-test; Figure 3A), and 3 of 3 donors having increased EC3 proportions upon ERG knock-down (Figure 3B).

**Figure 3.**
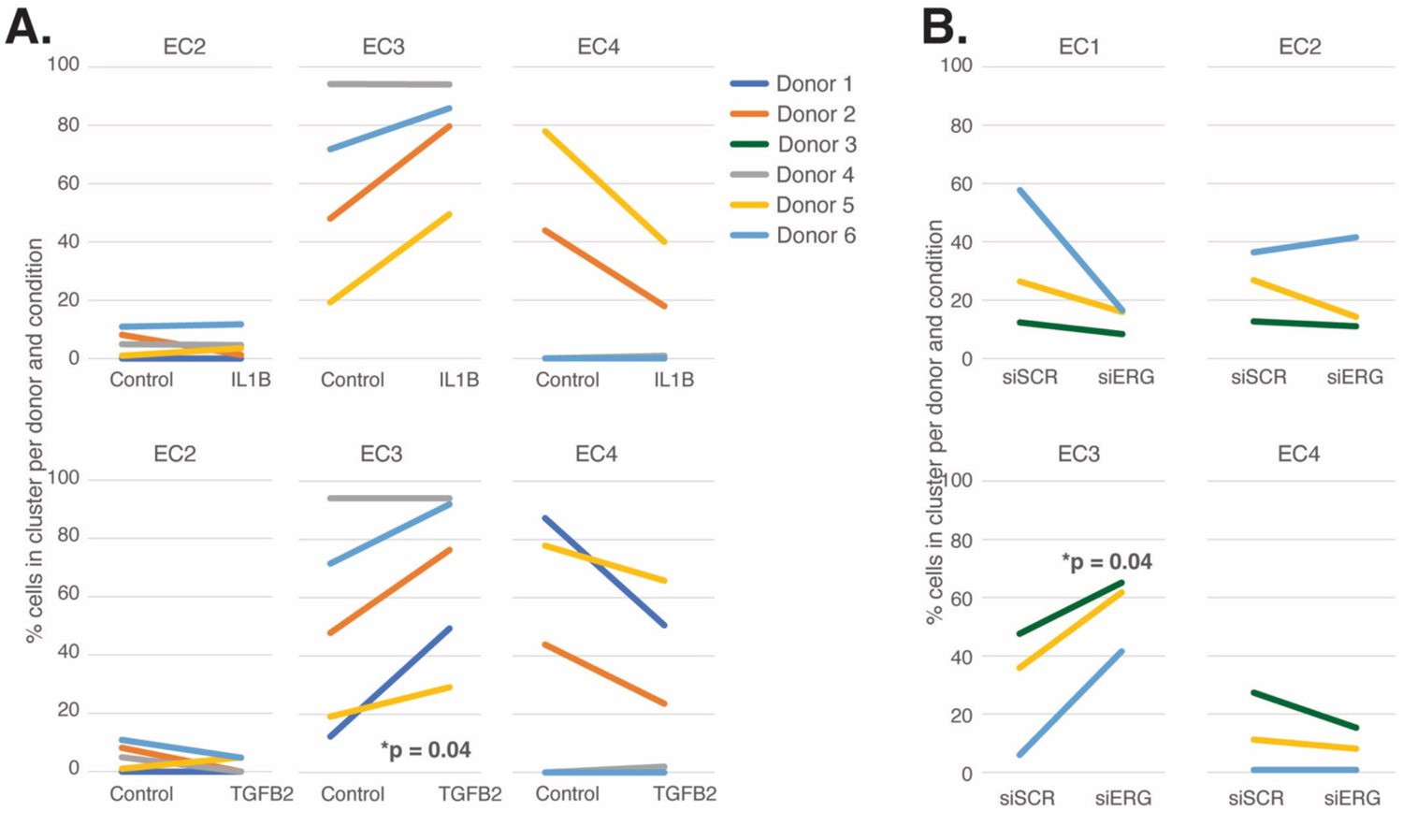
EC activating perturbations modestly shift cells into the EC3 subtype. **(A)**, The proportion of cells in 7-day control and 7-day IL1B treatment are shown per HAEC donor and cluster on the top and for 7-day control and 7-day TGFB2 on the bottom **(B)**, The proportion of cells in 7-day siSCR control and 7-day siERG knock-down are shown per HAEC donor and cluster. EC1 was omitted in A due to lack of cells in both conditions.

In addition to heterogeneity across EC clusters, data in Figure 3 underscores that there is heterogeneity among EC cultures. To quantify this effect, we performed principal component (PC) analysis to evaluate the overall contributions that donor and experimental conditions have on variance in this dataset. We found that pro-EndMT perturbations elicited greater variance in RNA expression (38-56% of variance) than donor (17%-27% variance) (**Figures S2A-C** in the **Data Supplement**), supporting that the transcriptional and epigenetic programs elicited by experimental conditions have a greater overall consequence than donor. This finding provides the opportunity to elucidate how different EC clusters respond to pro-EndMT exposures across genetically distinct ECs.

### Pro-EndMT Perturbations *In Vitro* Elicit EC Subtype-Specific Transcriptional Responses

We next sought to evaluate the similarities and differences among pro-EndMT perturbations and evaluate the transcriptional response elicited in each EC subtype. Differential gene expression analysis was performed using pseudo-bulked profiles grouped by donor, subcluster, and experimental groupings (**Table S9** in the **Data Supplement**).

Overall, we found heterogeneity in transcriptional responses across EC subtypes. While EC1 and EC2 transcripts were predominantly perturbed by siERG, the greatest number of transcripts differentially expressed in EC3 were those responsive to IL1B, though siERG and TGFB2 also regulated tens to hundreds of transcripts in EC3. In contrast, transcripts in EC4 were predominantly responsive to TGFB2 (Figure 4A, **Table S9** in the **Data Supplement**). With respect to EC4, we questioned whether transcripts were predominantly responsive to TGFB2 due to differences in expression of TGFB receptors. While we observed increased TGFBR1 expression in EC4, we observed relatively less expression of TGFBR2 and ACVRL1 in EC4 when compared to EC1, EC2, and EC3 (**Figure S3A** in the **Data Supplement**). We next questioned whether EC3 transcripts were predominantly responsive to IL1B due to differences in IL1B receptor expression. Notably, we did not observe differences in IL1B receptor expression, suggesting that their transcription is not responsible for divergent EC responses across EC subtypes (**Figure S3B** in the **Data Supplement**). Interestingly, we did observe differential expression of IL1RL1 in EC2, which may influence EC2 response to cytokine (**Figure S3B** in the **Data Supplement**).

**Figure 4.**
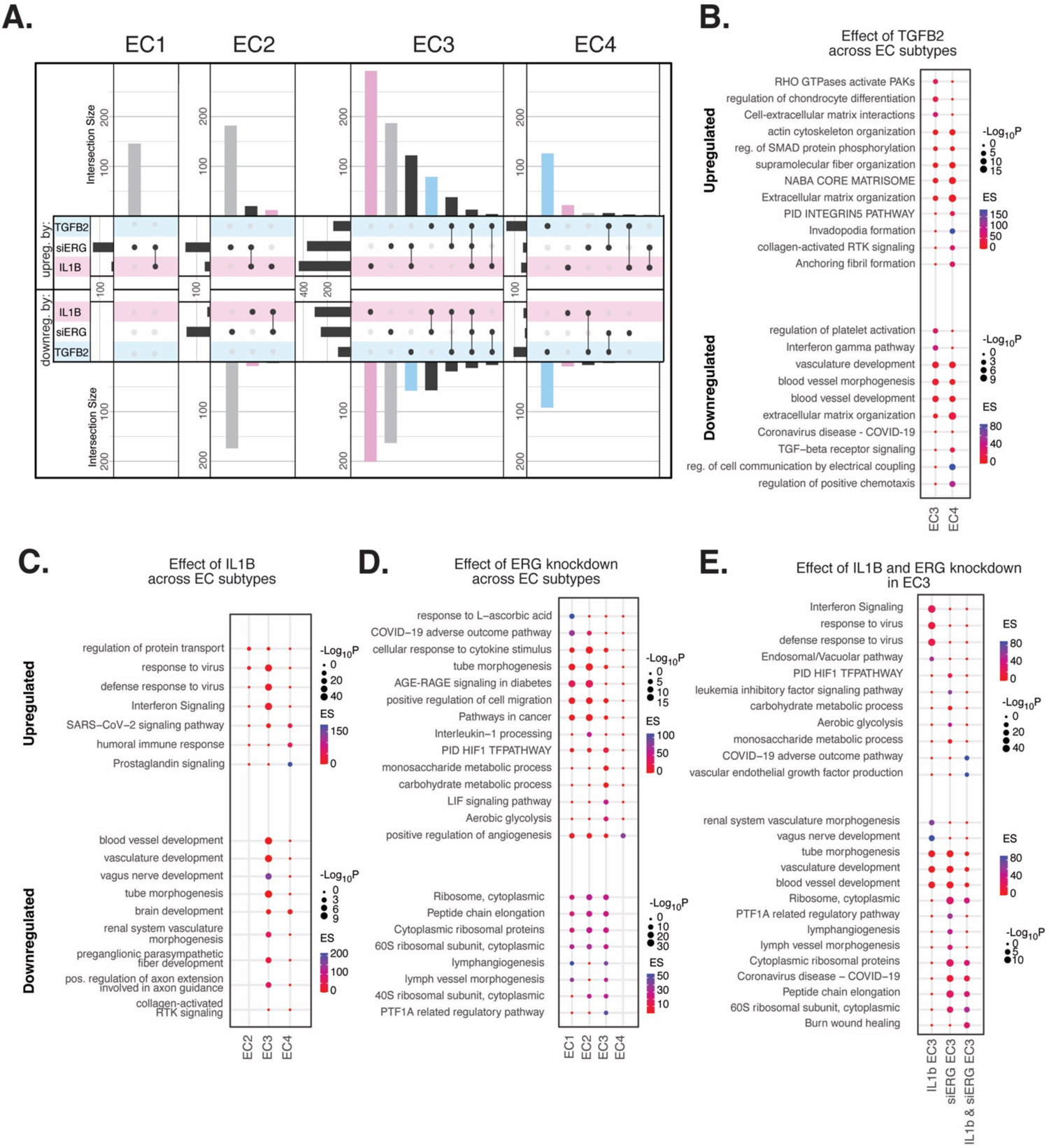
EC activating perturbations *in vitro* elicit EC subtype-specific transcriptional responses. **(A)**, Upset plots of up- and down-regulated DEGs across EC subtypes with siERG (grey), IL1B (pink), and TGFB2 (blue). Upset plots visualize intersections between sets in a matrix, where the columns of the matrix correspond to the sets, and the rows correspond to the intersections. Intersection size represents the number of genes at each intersection. **(B)**, PEA for EC3-4 up- and down-regulated DEGs with TGFB2 compared to control media. **(C)**, PEA for EC2-4 up- and down-regulated DEGs with IL1B compared to control media. **(D)**, PEA for EC1-4 up- and down-regulated DEGs with siERG compared to siSCR. **(E)**, PEA comparing up- and down-regulated DEGs that are mutually exclusive and shared between IL1B and siERG in EC3.

When comparing enriched pathways across perturbations, we observed that over 80% of transcripts differentially expressed by a treatment in EC4 were in response to TGFB2 (Figure 4A, **Table S9** in the **Data Supplement**). TGFB2-affected transcripts for EC4 were enriched in invadopodia formation (R-HAS-8941237; p-value 2.7×10^-7^) and anchoring fibril formation (R-HAS-2214320; p-value 3.6×10^-7^) (Figure 4B). Notably, TGFB2-affected genes for EC3 share several mesenchymal-related enriched pathways with TGFB2-affected genes for EC4, including actin cytoskeleton organization (GO:0030036; p-value 4.4×10^-7^), NABA CORE MATRISOME (M5884; p-value 2.8×10^-7^), and ECM organization (R-HSA-1474244; p-value 5.4×10^-7^). TGFB2-attenuated transcripts unique to EC3 were enriched in platelet activation (GO:0030168; p-value 1.4×10^-4^) (Figure 4B).

Most transcripts affected in EC3 were responsive to IL1B (Figure 4A). Importantly, several EC3 genes differentially expressed with IL1B were also affected with siERG (Figure 4A). IL1B-affected transcripts in EC3 are not enriched in mesenchymal-like pathways (Figure 4C). However, EC3 IL1B-attenuated genes are enriched in blood vessel development (GO:0032502; p-value 5.1×10^-11^), indicating that this perturbation still has anti-endothelial effects (Figure 4C).

Most genes significantly affected by perturbations in EC1 and EC2 were responsive to siERG, likely due to their more endothelial-like phenotypes compared to EC3 and EC4 (Figure 4A). siERG-affected genes in EC1 and EC2 were enriched in COVID-19 adverse outcome pathway (52) (WP4891; p-values 5×10^-9^ and 8.3×10^-5^, for EC1-2 respectively) and AGE-RAGE signaling in diabetes (53) (hsa04933; p-values 8.9×10^-16^ and 1.9×10^-20^, respectively), while EC3 siERG-perturbed genes are enriched with a unique metabolic profile demonstrated by enrichment in monosaccharide metabolic process (GO:0005996; p-value 1×10^-6^), carbohydrate metabolic process (GO:0005975; p-value 6.6×10^-7^), and aerobic glycolysis (WP4629; p-value 4.1×10^-5^) (Figure 4D). In contrast, EC4 siERG-induced genes are enriched in positive regulation of angiogenesis (GO:0045766; p-value 4.5×10^-6^), a function normally impaired in *ERG*-depleted endothelial cells (Figure 4D) (38).

Due to the role that ERG plays in inhibiting NF-KB-dependent inflammation *in vitro* and *in vivo* (37), we set out to characterize mutually exclusive and shared pathways between IL1B and siERG (Figure 4E). Importantly, siERG, but not IL1B-perturbed genes, involve several previously mentioned metabolic processes including carbohydrate metabolic process (GO:0005975; p-value 6.6×10^-7^), aerobic glycolysis (WP4629; p-value 4.1×10^-5^), and monosaccharide metabolic process (GO:0005996; p-value 1×10^-6^). This suggests differences in the ability of ERG and IL1B to modify metabolism. Interestingly, IL1B but not siERG upregulated interferon signaling and viral responsive pathways (GO:0051607, p-value 1×10^-37^; R-HSA-913531, p-value 1×10^-41^). Shared IL1B- and siERG-upregulated genes were enriched in COVID-19 adverse outcome pathway (WP4891; p-value 1.9×10^-9^) (52). Shared IL1B- and siERG-attenuated genes are enriched in several processes involving ribosomal proteins, including ribosome, cytoplasmic (CORUM:306; p-value 3.3×10^-7^), cytoplasmic ribosomal proteins (WP477; p-value 5.3×10^-7^), and peptide chain elongation (R-HSA-156902; p-value 5.9×10^-7^) (Figure 4E). This finding indicates that the downregulation of ribosomal genes is a hallmark of inflammatory and *ERG*-depleted endothelium. Altogether, these data demonstrate the heterogeneity in EC subtype response to pro-EndMT perturbations.

### *In Vitro* EC EndMT Models Reorganize Epigenetic Landscapes with Subtype Specificity

To gain insight into gene regulatory mechanisms responsible for EC subtype transcriptional responses to IL1B, TGFB2, and siERG, we compared the effects of these perturbations on chromatin accessibility. Across all three treatments, we identified 4,034 differentially accessible regions (DARs, **Table S10** in the **Data Supplement**, **Methods**). The majority of DARs for EC1 and EC2 were responsive to siERG, while the majority of DARs for EC3 were responsive to IL1B (**Figure S4A** in the **Data Supplement**, **Table S10** in the **Data Supplement**). Interestingly, the epigenetic landscape of EC4 differs from its transcriptional response, insofar as most peaks were responsive to IL1B (not TGFB2) (**Figure S4A** in the **Data Supplement**, **Table S10** in the **Data Supplement**). To inform the TFs likely bound to differentially accessible regulatory elements, motif enrichment analysis was performed (**Figure S4B-D** in the **Data Supplement**). Several distinct TF motifs were enriched across EC subtypes. For IL1B, we observed enrichment in KLF15 (adjusted p-value 5×10^-10^) (kruppel like factor 15) in EC2 alone (**Figure S4B** in the **Data Supplement**). siERG induced peaks showed subtype-specific motif enrichments, including TWIST1 (adjusted p-value 2.5×10^-22^) (twist family bHLH transcription factor 1), HAND2 (adjusted p-value 2.3×10^-19^) (heart and neural crest derivatives expressed 2) for EC1, RELA (adjusted p-value 9.5×10^-20^) (RELA proto-oncogene, NF-KB subunit) for EC2, and CEBPD (adjusted p-value 1.6×10^-29^) for EC3 (**Figure S4C** in the **Data Supplement**). Minimal motif enrichment was observed with siERG for EC4.

We also found several TF motifs enriched across more than one EC subtype upon perturbation. IL1B-affected peaks gained in EC1 and EC2 shared enrichments for TFDP1 (adjusted p-value 1.3×10^-4^ and 9×10^-4^ for EC1 and EC2, respectively) (transcription factor Dp1) and ZBTB14 motifs (adjusted p-value 2.2×10^-4^ and 2×10^-8^, respectively) (zinc finger and BTB domain containing 14). IL1B-induced peaks in EC3 and EC4 shared enrichment for CEBPD (adjusted p-value 4.4×10^-73^ and 1.6×10^-33^ for EC3 and EC4, respectively) and CEBPG motifs (adjusted p-value 5.4×10^-45^ and 7.1×10^-18^, respectively) (CCAAT enhancer binding protein delta and gamma) (**Figure S4B** in the **Data Supplement**). TGFB2-affected peaks in EC1, EC2, and EC3 shared enrichment for ZBTB14 (adjusted p-values 6.8×10^-31^, 5.1×10^-12^, and 2×10^-8^, for EC1, EC2, and EC3, respectively) while TGFB2-induced peaks in EC3 and EC4 shared enrichment for the SMAD5 motif (adjusted p-value 7.4×10^-6^ and 4.2×10^-11^, for EC3 and EC4, respectively) (SMAD family member 5) (**Figure S4D** in the **Data Supplement**). Taken together, while several enriched motifs are shared across EC subtypes, divergent epigenetic landscapes are also induced with pro-EndMT perturbations. We therefore conclude that different transcriptional responses to these perturbations across EC subtypes are elicited by distinct TFs, including members of families of the KLF, TWIST, HAND, p65, and CEBP families.

### Meta-Analysis of *Ex Vivo* Human Atherosclerotic Plaque snRNA-seq Datasets

To understand the diversity of ECs in human atherosclerotic plaques and evaluate their relationships to our *in vitro* system, we performed a meta-analysis of data from recent publications that utilized scRNA-seq from human atherosclerotic lesions (26–29) (accessions in **Table S11** in the **Data Supplement**). We identified 24 diverse clusters among 58,129 cells after integration of 17 different coronary and carotid samples (Figure 5A and **Table S12** in the **Data Supplement**). Clusters were annotated using a combinatorial approach including canonical marker genes, CIPR (54), and the original publications (Figure 5B). Clusters were annotated as: T-lymphocytes, natural killer T-cells, ECs, macrophages, VSMCs, fibroblasts, B-lymphocytes, basophils, neurons, and plasmacytoid dendritic cells (PDCs) (Figure 5A). We find the greatest proportion of cells belonging to each major cell type derive from carotid arteries, except for neurons which derive exclusively from coronary arteries, and PDCs which derive exclusively from carotid arteries (**Figures S5B-C** in the **Data Supplement**). Expected pathway enrichments are observed for annotated cell types, including NABA CORE MATRISOME (M5884; p-value 4.8×10^-41^) for fibroblasts, blood vessel development (GO:0001568; p-value 5.6×10^-33^) for ECs, and actin cytoskeleton organization (GO:0030036; p-value 1.3×10^-15^) for VSMCs (**Figure S5D**-**G** in the **Data Supplement**). These observations support the diverse composition of human atherosclerotic lesions.

**Figure 5.**
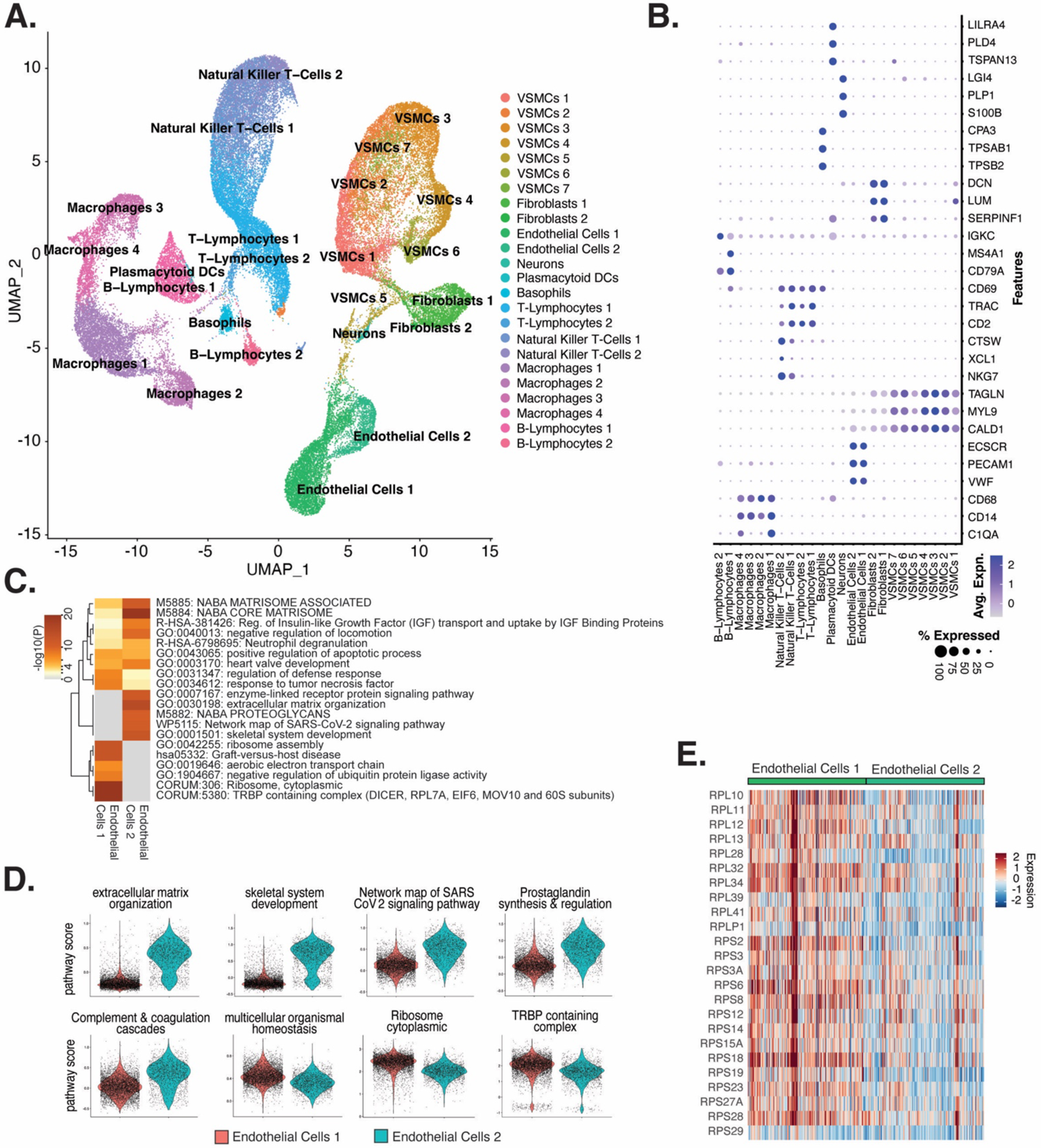
ECs from *ex vivo* human atherosclerotic plaques show two major populations. **(A)**, scRNA-seq UMAP of different cell subtypes across 17 samples of *ex vivo* human atherosclerotic plaques. **(B)**, Dot plot of top markers for each cell type. **(C)**, Heatmap of pathway enrichment analysis (PEA) results generated from submitting 200 differentially expressed genes (DEGs) between Endothelial Cells 1 (Endo1) and Endothelial Cells 2 (Endo2). Rows (pathways) and columns (cell subtypes) are clustered based on -Log_10_(P)**. (E)**, Heatmap displaying expression of genes belonging to ribosome cytoplasmic pathway for Endo1 and Endo2.

We evaluated what pathways distinguished the Endothelial Cells 1 (Endo1) and Endothelial Cells 2 (Endo2) subtypes from our *ex vivo* meta-analysis (Figure 5C). We found Endo2 has an EndMT-related phenotype, with enrichment in mesenchymal pathways including NABA MATRISOME ASSOCIATED (M5885; p-value 1.6×10^-14^), ECM organization (R-HSA-1474244; p-value 6×10^-17^), skeletal system development (GO:0001501; p-value 3.4×10^-13^), and network map of SARS-CoV-2 signaling pathway (52) (WP5115; p-value 1.3×10^-11^) (Figure 5C-D). Additionally, we observe enrichment for inflammatory pathways in Endo2 including prostaglandin synthesis and regulation (WP98; p-value 1.2×10^-7^), and complement and coagulation cascades (hsa04610; 1×10^-10^) (Figure 5C-D) (55, 56). On the contrary, Endo1 was highly enriched in multicellular organismal homeostasis (GO:0048871; p-value 3.3×10^-8^) and lowly enriched in mesenchymal pathways (M5885; p-value 1×10^-3^; no enrichment for R-HSA-1474244, GO:0001501, or WP5115), indicating a canonical EC phenotype (Figure 5C-D). Interestingly, Endo1, but not Endo2, is highly enriched in ribosome, cytoplasmic pathway (CORUM:306; p-value 9.3×10^-96^), and TRBP containing complex (CORUM:5380; DICER, RPL7A, EIF6, MOV10 and subunits of the 60S ribosomal particle; p-value 1.5×10^-22^), suggesting a potential protective role for this complex along with ribosomal gene expression (57, 58). The depletion of these pathways may serve as a hallmark of activated endothelium (Figure 5C-E). We interpret these results to suggest that Endo1 is a classical endothelial state, while Endo2 appears to be characterized by ECM production and possibly indicate EndMT. This interpretation is further corroborated by evidence of upregulation of several classical EndMT markers in Endo2, including: *FN1*, *BGN*, *COL8A1*, *ELN*, *CCN1*, and *FBLN5* (**Figure S6** in the **Data Supplement**) (59–64).

### *Ex Vivo*-derived Module Score Analysis Reveals Differences among *In Vitro* EC Subtypes and EndMT Stimuli

To directly evaluate relationships between the *ex vivo* and *in vitro* cell subpopulations, we utilized module scores. These quantitative scores are based on the sum of *ex vivo* marker genes across each cluster and were used to evaluate similarity to each *in vitro* cell subcluster. Unexpectedly, the *ex vivo* cluster that consistently generated the greatest module scores for *in vitro* ECs is the VSMCs cluster 5 (VSMC5) (Figure 5A; **Figure S7A** in the **Data Supplement**). VSMC5 bridges the EC to SMC and fibroblast clusters in the *ex vivo* analysis (Figure 5A). Marker genes of VSMC5 are expressed across *ex vivo* and *in vitro* clusters (**Figure S8A** in the **Data Supplement**) and include important regulators of ECM, such as *BGN*, *VCAN*, *FN1*, as well as several collagen genes (*COL1A1*, *COL1A2*, *COL3A1*, *COL6A1*) (**Figure S8A-B** in the **Data Supplement**). VSMC5 marker transcripts also include several lncRNAs and mitochondrial transcripts (*CARMN*, *MALAT1*, *NEAT1*; *MT-ATP6*, *MT-ND4*, and *MT-CYB*) (**Figure S8A** in the **Data Supplement**). *Ex vivo* Endo1 and Endo2 module scores are the second highest scoring across *in vitro* clusters. Cells scoring high for Endo1 are concentrated in the *in vitro* EC1 cluster, while cells scoring high in Endo2 are concentrated to the *in vitro* EC3 locale (**Figure S7B-E** in the **Data Supplement**). This supports that EC3 is a more activated subtype than EC1. Finally, among *in vitro* cells, those with highest VSMC5 module scores are concentrated in EC4, underscoring that EC4 is a more mesenchymal sub-phenotype *in vitro* (**Figure S7B-E** in the **Data Supplement**).

We stratify these analyses by pro-EndMT treatment and find greater VSMC5 module scores with TGFB2 treatment versus control for EC3 (adjusted p-value = 0.001) and EC4 (adjusted p-value = 9.9×10^-15^) (**Figure S9A-C** in the **Data Supplement**). However, there is no difference in VSMC5 module scores for EC1-2 between control and TGFB2 treatment, suggesting these subtypes are resistant to transcriptional changes by TGFB2 exposure (i.e., EC1-2). This is in contract to the more mesenchymal-like EC (i.e., EC3-4) subtypes which are more responsive to TGFB2 (**Figure S9A-C** and **Table S12**-**13** in the **Data Supplement**). We observe siERG lowers Endo1 scores across all EC subtypes (adjusted p=9.9×10^-15^ for EC1-4), indicating *ERG* depletion decreases endothelial-likeness across all EC subtypes (**Figure S9A-C** and **Table S13**-**14** in the **Data Supplement**). Moreover, siERG increases VSMC5 scores for EC2 (adjusted p=2.8×10^-9^) and EC3 (adjusted p-value 0.04), indicating siERG elicits activated and mesenchymal characteristics (**Figure S9A-C** and **Table S13**-**14** in the **Data Supplement**).

### EC Subtype is a Major Determinant in Modeling Cell States Observed in Atherosclerosis

In addition to module score analysis, we applied a complementary approach to quantitatively relate *in vitro* EC subtypes and pro-EndMT perturbations to *ex vivo* cell types. We calculate average expression profiles for all major cell populations in both *ex vivo* and *in vitro* datasets and examine the comprehensive pairwise relationship among populations with hierarchical clustering using Spearman Correlation (Figure 6A). All *in vitro* transcripts significantly regulated across all pro-EndMT perturbations at 5% False Discovery Rate (FDR) (65) are used in this analysis, although several additional means to select transcripts showed similar results. This analysis reveals three major observations. First, *in vitro* EC4 cells are most like mesenchymal *ex vivo* cell types including VSMCs and fibroblasts (indicated by the yellow block of correlations in the bottom left of the heatmap in Figure 6A). Second, *in vitro* EC1, EC2, and EC3 are most like *ex vivo* Endo1 and Endo2 populations, especially among the siSCR and 7-day control cells. Moreover, cells in the siSCR condition in EC1 are most like *ex vivo* Endo1, reinforcing that these two populations are the most canonically ‘healthy’ endothelial populations. Third, while pro-EndMT perturbations did elicit variation in how similar *in vitro* ECs resembled *ex vivo* transcriptomic signatures, these effects are secondary to which subtype the cells belonged (Figure 6A). Taken together, these findings underscore that EC subtype, versus perturbation, is a greater determinant of similarity to *ex vivo* cell types.

**Figure 6.**
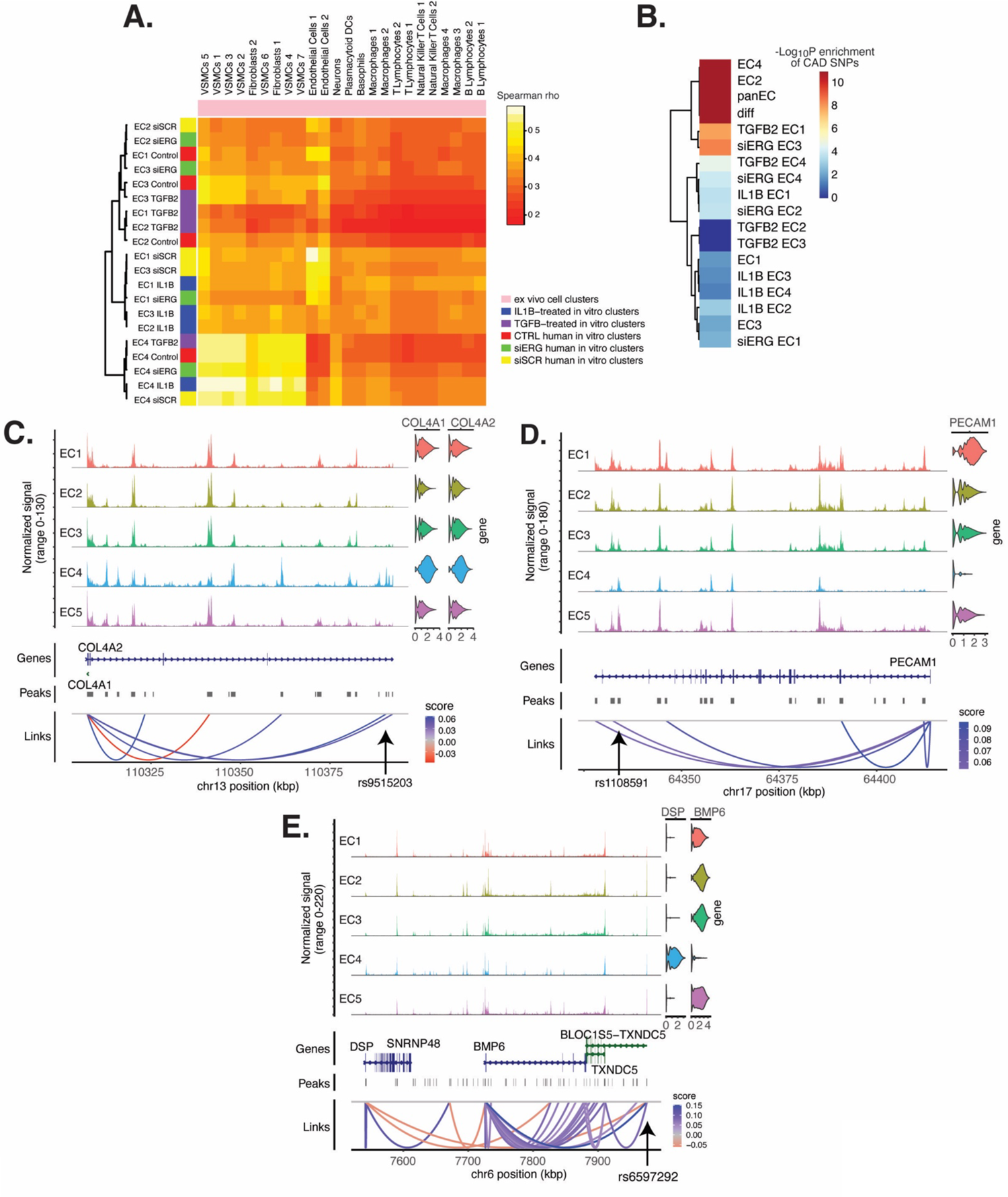
EC subtype is a major determinant in the ability to recapitulate ‘omic profiles seen in atherosclerosis. **(A)**, Heatmap displaying average expression between *in vitro* perturbation-subtype combinations and *ex vivo* cell subtypes using all up- and down-regulated genes between IL1B, TGFB2, or siERG versus respective controls. Spearman correlation was used as the distance metric. Rows (*in vitro* EC subtypes) and columns (*ex vivo* cell subtypes) are clustered using all significant genes (adjusted p-value < 0.05) induced and attenuated across all *in vitro* EC subtypes for each perturbation versus its respective control. **(B)**, Heatmap of CAD-associated SNP enrichments across *in vitro* EC subtypes and perturbation-subtype combinations. Rows (EC subtypes and perturbation-subtype combinations) are clustered using -Log_10_(P) for enrichment in significant CAD-associated SNPs (p-value < 5×10^-8^). Note that “diff” represents peaks common to more than one EC subtype; it is found by subtracting EC1-5 subtype-specific peaks from the entire *in vitro* peak set (termed “panEC”). **(C)**, Coverage plots displaying links for *COL4A1*/*COL4A2* genes to EC4-specific peaks, including one overlapping with CAD-associated SNP rs9515203. **(D)**, Coverage plot showing links for *PECAM1* gene to EC4-specific peaks, including one overlapping with CAD-associated SNP rs1108591. **(E)**, Coverage plot showing links for *BMP6* gene to EC4-specific peaks, including one overlapping with CAD-associated SNP rs6597292.

### CAD-Associated Genetic Variants Are Enriched Across EC Subtype Epigenomes

Genetic predisposition to CAD is approximately 50% heritable with hundreds to thousands of genetic loci supposed to be involved in shaping an individual’s propensity for disease (66, 67). Most CAD-associated variants are not protein coding, suggesting they perturb cellular function through gene regulatory functions.

We therefore asked whether the open chromatin regions in this *in vitro* dataset coincided with locations of single nucleotide polymorphisms (SNPs) reported in the latest CAD meta-GWAS analysis from the Millions Veterans Project (MVP), which includes datasets from CARDIoGRAMplusC4D 1000G study, UK Biobank CAD study, and Biobank Japan (6). We found significant enrichment in CAD-associated SNPs for the complete set of accessible regions across all EC subtypes (termed “panEC”; adjusted p-value 1.5e×10^-93^; Odds Ratio (OR)=1.8; Figure 6B, **Table S15-16** in the **Data Supplement**) when comparing CAD SNPs exceeding the genome-wide significance threshold of p<5×10^-8^ versus non-significant SNPs (**Methods**). Among accessible regions unique to EC subtypes, EC4 shows the greatest enrichment (adjusted p-value 7.85×10^-6^; OR=1.74). Additionally, EC2 is also enriched for CAD SNPs (adjusted p-value 6.3×10^-8^; OR=2.15), supporting a role for proliferative ECs in CAD. Of all accessible regions influenced by pro-EndMT perturbations, siERG and TGFB2 sets are most enriched for CAD variants (Figure 6B**, Table S15**-**16** in the **Data Supplement**).

The measurement of both gene expression and DNA accessibility in the same cell enables testing for direct correlation, or ‘links’, between accessibility of noncoding DNA elements and gene expression of their potential regulatory targets (i.e., gene promoters). This is achieved by testing for correlation between DNA accessibility and the expression of a nearby gene across single cells (48, 68). Focusing on EC4, we search for EC4-specific sites of correlated chromatin accessibility and linked target gene expression. Upon restricting linked peaks overlapping CAD SNPs, we identify 81 significant SNP-peak-gene trios (p < 0.05) representing 46 unique genes with specific activity in EC4 (**Table S17** in the **Data Supplement**). We submit the 46 unique genes to Metascape (69) and observe enrichment in EndMT-related pathways including blood vessel development (GO:0001568; p-value 2.1×10^-10^), crosslinking of collagen fibrils (R-HSA-2243919; p-value 1.4×10^-8^), and canonical and non-canonical TGF-B signaling (WP3874; p-value 2.2×10^-6^) (**Figure S10** in the **Data Supplement**). Literature review of this gene list further confirms several linked EC4-restricted genes associated with cardiovascular disease, including *COL4A1*, *COL4A2*, *PECAM1, DSP*, and *BMP6*, (Figure 6C-E) (70–72). Altogether, these data underscore that common genetic variation influences individual propensities for CAD through ECM-organizing functions evidenced by the EC4 phenotype.

## DISCUSSION

The major goals of this study were fourfold: (1) to quantitatively assess molecular heterogeneity of cultured HAECs *in vitro*, (2) to evaluate and compare molecular changes elicited by EC activating perturbations at single cell resolution, (3) to assess similarities between *in vitro* and *ex vivo* EC signatures to inform the extent to which *in vitro* models recapitulate *ex vivo* biology, and (4) investigate how heterogeneous EC populations are enriched for genetic associations to CAD. Findings for each of these goals are discussed below along with important implications and questions arising from this work.

The multiomic single cell profiles of 15,220 cells cultured *in vitro* from six individuals enabled the discovery of 5 EC subpopulations, named EC1, EC2, EC3, EC4, and EC5. Except for EC5, EC subpopulations were comprised of cells from multiple donors and perturbations, which lends credence to the reproducibility of these biological states. The loosely defined phenotypes, based on pathway enrichment analysis, were healthy/angiogenic for EC1, proliferative for EC2, activated for EC3, and mesenchymal for EC4. Angiogenic (9, 10, 14), proliferative (19, 73), and mesenchymal (19) ECs have been previously reported in literature. The three activating perturbations (TGFB2, IL1B, siERG) had markedly unique effects on different EC subclusters, highlighting the fact that *in vitro* systems contain populations of discrete cell subtypes, or states, that respond divergently to even reductionistic experimental conditions. Implications of such heterogeneity include both a need to elucidate what factors dictate treatment responsiveness, as well as experimental design and data interpretation that considers heterogeneity of response. The exact origin of EC heterogeneity observed in this study is unknown. We consider it likely that EC1 EC2, EC3, and EC4 subpopulations, which were populated by most donors, date back to the original isolation of ECs from aortic trimmings, implying that different states were preserved across passage in the culture conditions. However, we cannot exclude the possibility that some of the subpopulations have expanded since seeding of the cultures. If that were the case, EC1, EC2, EC3, and EC4 represent reproducible cell states consequent to primary culture of arterial cells. In fact, the limited correlation with *ex vivo* data supports this interpretation. Future studies will be required to delineate the exact source of heterogeneity in these systems.

In this study, we set out to elucidate whether the mesenchymal phenotype of EC4 was an end-stage result of EndMT and whether TGFB2, IL1B, and/or siERG would increase the proportion of cells in EC4. As shown in Figure 3, this hypothesis was incorrect, and the only cluster with a modest increase in cell proportions upon stimulation was EC3. Moreover, while the percent of cells in EC3 increased with TGFB or IL1B, they decreased in EC4, suggesting trans-differentiation from EC4 into EC3 with these perturbations. We cannot exclude the possibility that EC3 is an EndMT cluster, although we would have expected more significant deviation from clusters EC1 and EC2. It is also possible that the postmortem state experienced by aortic explants prioir to EC isolation could induce changes in the ECs, or that the duration and doses of perturbations chosen were not sufficient to elicit complete EndMT. While the duration and doses employed in our study were established based on literature reports reporting EndMT phenotypes (33, 50, 74), EndMT was quantified by expression of only a few marker genes rather than complete transcriptomic analysis. This raises an important conclusion of our study, which is that EndMT is not well-defined molecularly and it remains possible that several different molecular profiles may each represent variant flavors of EndMT.

We found that TGFB2, IL1B, and siERG have many distinct effects on EC molecular profiles (Figures 3-4). In general, TGFB2 elicits a greater transcriptomic and epigenomic response in the mesenchymal EC subtype, EC4, while siERG and IL1B regulate the greatest numbers of shared transcripts and chromatin regions in more endothelial clusters EC1, EC2, and EC3. One interpretation for this finding is that IL1B treatment and depletion of *ERG* directly affect rewiring transcription in ECs while TGFB2 may affect other cell types in the vascular wall (or culture plate) that in turn affect ECs through paracrine interactions. Part of the similarities between IL1B and siERG responses may be explained by the fact that *ERG* depletion increases IL1B production (41).

A major question raised by this work is the origin of cells in the mesenchymal cluster EC4. We originally hypothesized this cluster was the result of EndMT, which led to our investigations as to whether we could leverage EndMT-promoting exposures (IL1B, TGFB2, siERG) *in vitro* observe an expansion of treated cells in the EC4 population. To our surprise, the EC4 population did not expand. If anything, these exposures reduced the proportion of cells in ECs (Figure 4). Nonetheless, it remains a possibility that EC4 represents cells that had undergone EndMT *in vivo* prior to culture and that the exposures we presented *in vitro* were not sufficient to elicit a complete EndMT transition. Another viable hypothesis is that cells in EC4 are of SMC origin and have persisted in culture alongside their EC counterparts. Cells used in this study were isolated by luminal collagenase digestion of explanted aortic segments and were tested at early passage for EC phenotypic markers including VWF expression, cobblestone morphology, and uptake of acetylated LDL. Notably, these rigorous metrics to ensure pure EC isolation occurred prior to our group’s studies. In addition, if some of the isolated cells had undergone EndMT *in vivo* prior to isolation, it would be nearly impossible to distinguish their cell of origin after isolation since their collective molecular phenotypes would appear as an SMC. Without lineage tracing, which is currently not possible in human tissue explants, it would not be possible to distinguish cell origin. Nonetheless, this remains an important issue that is the subject of ongoing investigations. What we can confidently discern from these data is that these distinct cell sub-populations respond differently to the disease-relevant exposures of IL1B, TGFB2, and ERG depletion.

The current study sought to evaluate similarities and differences between *in vitro* primary cultures of HAECs to *ex vivo* single cell signatures of cells from human lesions. First, we leveraged transcriptomic profiles from clusters in the scRNA meta-analysis of human lesions and evaluated each *in vitro* cluster using a module score (Figures 5 and **Figure S8** in the **Data Supplement**). The three *ex vivo* clusters with greatest similarity to *in vitro* clusters were Endo1, Endo2, and VSMC5. Pathway enrichment analysis suggested that the *ex vivo* Endo1 cluster is close to the classic “healthy” EC state relative to Endo2, which returned pathway enrichments consistent with activated endothelium (Figure 5C-D). Interestingly, Endo2 is depleted in ribosome transcripts as well as transcripts in the Dicer complex (Figure 5C-E), which may serve as hallmarks of dysregulated endothelium *in vivo*. VSMC5 is an interesting *ex vivo* cluster insofar as it spans the endothelial, fibroblast, and VSMC clusters (Figure 5A) and is enriched for genes in actin cytoskeleton, extracellular matrix organization, and more (**Figure S8** in the **Data Supplement**). *In vitro* EC1, EC2, and EC3 score generally greater in Endo1 and Endo2 relative to the more mesenchymal EC4 (**Figure S7** in the **Data Supplement**). Consistent with the intent of the pro-EndMT treatments, they generally decrease Endo1 and Endo2 scores and increase VSMC5 scores. However, these effects are unexceptional in comparison to effects of EC subtype. In addition to module scores, we also utilized unsupervised clustering of Spearman correlation coefficients across *ex vivo* and *in vitro* average gene expression profiles, finding again that EC1, EC2, and EC3 are more like Endo1 and Endo2 and EC4 is more like VSMCs (Figure 6A). As expected, the control (siSCR) cells are most correlated to healthy Endo1 transcriptomes; however, the correlation coefficient achieved is modest, at rho = 0.56. We cannot exclude the possibility that the moderate correlation coefficient observed between *in vitro* and *ex vivo* ECs may be explained by anatomic differences (i.e., aortic versus coronary and carotid arteries). While reinforcing that *in vitro* cell cultures best resemble ECs isolated *ex vivo*, regardless of perturbation, this finding accentuates how different cultured cells are and paves the way for quantitatively evaluating and improving *in vitro* models.

Finally, GWAS studies have established that hundreds of independent common genetic variants in human populations affect risk for CAD, yet discovering the causal mechanisms remains a major challenge given that most of the risk is in non-coding regions of the genome. One approach to prioritize causal variants in regulatory elements is through integration of open chromatin regions from the cell type and states of interest followed by expression quantitative trait loci (eQTL) or other linking evidence to target gene (75, 76). In the current study, we find significant enrichment for CAD-risk variants in open chromatin regions across the entire dataset (“panEC”) as well as specifically for EC2 and EC4 subpopulations (Figure 6B**; Table S15-17** in the **Data Supplement**). While EC3 was found to be more sensitive to perturbations in our *in vitro* experiments, we did not expect to see CAD-related SNPs enriched in EC3 because plasticity does not necessarily imply a pathological process. Moreover, while EC3 and EC4 both have mesenchymal phenotypes, EC3 may represent a reversible state that is lacking in EC4. This hypothesis would explain the enrichment of EC4, but not EC3, in CAD-related SNPs. Taken together, these data emphasize the value in multimodal datasets in human samples for prioritizing disease-associated SNPs and mechanisms.

## METHODS

### Tissue Procurement and Cell Culture

Primary HAECs were isolated from eight de-identified deceased heart donor aortic trimmings (belonging to three females and five males of Admixed Americans, European, and East Asian ancestries) at the University of California Los Angeles Hospital as described previously (42) (**Table S7** in the **Data Supplement**). The only clinically relevant information collected for each donor was their genotype (**Methods, “Genotyping and Multiplexing Cell Barcodes for Donor Identification”**). HAECs were isolated from the luminal surface of the aortic trimmings using collagenase, and identified by Navab et al. using their typical cobblestone morphology, presence of Factor VIII-related antigen, and uptake of acetylated LDL labeled with 1,1’-dioctadecyl-1-3,3,3’,3’-tetramethyl-indo-carbocyan-ine perchlorate (Di-acyetl-LD) (42). Cells were grown in culture in M-199 (ThermoFisher Scientific, Waltham, MA, MT-10-060-CV) supplemented with 1.2% sodium pyruvate (ThermoFisher Scientific, cat. no. 11360070), 1% 100X Pen Strep Glutamine (ThermoFisher Scientific, cat. no. 10378016), 20% fetal bovine serum (FBS, GE Healthcare, Hyclone, Pittsburgh, PA), 1.6% Endothelial Cell Growth Serum (Corning, Corning, NY, cat. no. 356006), 1.6% heparin, and 10μL/50 mL Amphotericin B (ThermoFisher Scientific, cat. no. 15290018). HAECs at low passage (passage 3-6) were treated prior to harvest every 2 days for 7 days with either 10 ng/mL TGFB2 (ThermoFisher Scientific, cat. no. 302B2002CF), IL1B (ThermoFisher Scientific, cat. no. 201LB005CF), or no additional protein, or two doses of small interfering RNA for ERG locus (siERG; **Table S18** in the **Data Supplement**), or randomized siRNA (siSCR; **Table S18** in the **Data Supplement**). Donors 7 and 8 were treated prior to harvest for 6 hours with either 1 ng/mL IL1B, or no additional protein, and included in the dataset during integration to generate the original UMAP (Figure 1B), but not used for the purposes of downstream analyses in this study (**Table S7** in the **Data Supplement**).

### siRNA Knock-down, qPCR, and Western Blotting

Knockdown of ERG was performed as previously described (41) using 1 nM siRNA oligonucleotides in OptiMEM (ThermoFisher Scientific, cat. no. 11058021) with Lipofectamine 2000 (ThermoFisher Scientific, cat. no. 11668030). Transfections were performed in serum-free media for 4 hours, then cells were grown in full growth media for 48 hours. All siRNAs and qPCR primers used in this study are listed in **Table S18** in the **Data Supplement**. Transfection efficiency for the siRNAs utilized in this study was verified using qPCR 7 days after transfection (**Figure S11A** in the **Data Supplement**). Protein knockdown is shown 2 days after transfection using the same siRNAs from a representative experiment (**Figure S11B** in the **Data Supplement**). Antibodies used included 1:1,000 recombinant anti-ERG antibody (ab133264) and 1:5,000 anti-histone H3 antibody (ab1791) (Abcam). Western blots were quantified using ImageJ (77).

### Nuclear Dissociation and Library Preparation

Nuclei from primary cells were isolated according to 10x Genomics *Nuclei Isolation for Single Cell Multiome ATAC + Gene Expression Sequencing* Demonstrated Protocol (CG000365, Rev C) (78). Nuclei were pooled isolated with lysis buffer consisting of 10 mM Tris-HCl (pH 7.5, Invitrogen, cat. no. 15567027), 10 mM NaCl (Invitrogen, cat. no. AM9759), 3 mM MgCl_2_ (Alfa Aesar, cat. no. J61014), 0.1% Tween-20 (ThermoFisher Scientific, cat. no. 9005-64-5), 0.1% IGEPAL CA-630 (ThermoFisher Scientific, cat. no. J61055.AP), 0.01% Digitonin (ThermoFisher Scientific, cat. no. BN2006), 1% BSA (Sigma Aldrich, cat. no. A2153), 1 mM DTT (ThermoFisher Scientific, cat. no. 707265ML), 1 U/μl RNase inhibitor (Sigma Protector RNase inhibitor; cat. no. 3335402001), and nuclease-free water (Invitrogen, cat. no. 10977015). The seven pooled samples were incubated on ice for 6.5 minutes with 100 μl lysis buffer and washed three times with 1 mL wash buffer consisting of 10 mM Tris-HCl, 10 mM NaCl, 3 mM MgCl_2_, 1% BSA, 0.1% Tween-20, 1 mM DTT, 1U/μl RNase inhibitor, and nuclease-free water. Samples were centrifuged at 500 rcf for 5 minutes at 4C, and the pellets were resuspended in chilled Diluted Nuclei Buffer consisting of 1X Nuclei Buffer (20X) (10X Genomics), 1 mM DTT (ThermoFisher Scientific, cat. no. 707265ML), 1 U/μl RNase inhibitor, and nuclease-free water. The homogenate was filtered through a 40-μm cell strainer (Flowmi, cat. no. BAH136800040) prior to proceeding immediately to 10X Chromium library preparation according to manufacturer protocol (CG000338).

### Genotyping and Multiplexing Cell Barcodes for Donor Identification

Genotyping of HAEC donors was performed as described previously (75). Briefly, IMPUTE2 (79) was used to impute genotypes utilizing all populations from the 1000 Genomes Project reference panel (phase 3) (80). Genotypes were called for imputed SNPs with allelic R2 values greater than 0.9. Mapping between genomic coordinates was performed using liftOver (81). VCF files were subset by genotypes for the donors of interest using VCFtools (82).

To identify donors across the *in vitro* dataset, snATAC- and snRNA-seq output BAM files from Cell Ranger ARC (10X Genomics, v.2.0.0) (43) were concatenated, sorted, and indexed using samtools (83). The concatenated BAM files were input with the genotype VCF file to demuxlet (84) to identify best matched donors for each cell barcode, using options “–field GT”. Verification of accurate donor identification was confirmed by visualizing female sex specific *XIST* for the known donor sexes (**Figure S12** in the **Data Supplement**).

### snRNA-seq Bioinformatics Workflow

A target of 10,000 nuclei were loaded onto each lane. Libraries were sequenced on NovaSeq6000. Reads were aligned to the GRCh38 (hg38) reference genome and quantified using Cell Ranger ARC (10X Genomics, v.2.0.0) (43). Datasets were subsequently preprocessed for RNA individually with Seurat version 4.3.0 (44). Seurat objects were created from each dataset, and cells with < 500 counts were removed. This is a quality control step, as it is thought that cells with low number of counts are poor data quality. Similarly, for each cell, the percentage of counts that come from mitochondrial genes was determined. Cells with > 20% mitochondrial gene percent expression (which are thought to be of low quality, possibly due to membrane rupture) were excluded. Demuxlet (84) was next used to remove doublets. The filtered library was subset and merged by pro-EndMT perturbation. Data were normalized with NormalizeData, and cell cycle regression was performed by generating cell cycle phase scores for each cell using CellCycleScoring, followed by regression of these using ScaleData (85). Batch effects by treatment were corrected using FindIntegrationAnchors using 10,000 anchors, followed by IntegrateData.

### snATAC-seq Bioinformatics Workflow

A target of 10,000 nuclei were loaded onto each lane. Libraries were sequenced on an NovaSeq 6000 according to manufacturer’s specifications at the University of Chicago. Reads were aligned to the GRCh38 (hg38) reference genome and quantified using Cell Ranger ARC (10X Genomics, v.2.0.0) (43). Datasets were subsequently preprocessed for ATAC individually with Seurat v4.3.0 (44) and Signac v1.6.0 (86) to remove low-quality nuclei (nucleosome signal > 2, transcription start site enrichment < 1, ATAC count < 500, and % mitochondrial genes > 20) (44). Next, demuxlet (84) was used to remove doublets. A common peak set was quantified across snATAC-seq libraries using FeatureMatrix, prior to merging each lane. A series of two iterative peak calling steps were performed. The first step consisted of calling peaks for every EndMT perturbation, and the second involved calling peaks for every cluster generated from Weighted Nearest Neighbor Analysis (WNN) (**Methods**, **“Integration and Weighted Nearest Neighbor Analyses”**). Latent semantic indexing (LSI) was computed after each iterative peak calling step using Signac standard workflow (48). Batch effects by treatment were finally corrected using FindIntegrationAnchors using 10,000 anchors, followed by IntegrateData.

### Integration and Weighted Nearest Neighbor Analyses

Following snRNA-seq and snATAC-seq quality control filtering, barcodes for each modality were matched, and both datasets were combined by adding the snATAC-seq assay and integrated LSI to the snRNA-seq assay. WNN (44) was next calculated on the combined dataset, followed by joint UMAP (_WNN_UMAP) visualization using Signac (48) functions FindMultimodalNeighbors, RunUMAP, and FindClusters, respectively. WNN is an unsupervised framework to learn the relative utility of each data type in each cell, enabling an integrative analysis of multimodal datasets. This process involves learning cell-specific modality “weights” and constructing a _WNN_UMAP that integrates the modalities. The subtypes discovered in the first round of WNN were utilized in an additional peak calling step for snATAC-seq, followed by latent semantic indexing (LSI) computation, re-integration, and a final round of WNN to achieve optimal peak predictions (**Methods, “Single Nucleus ATAC Sequencing Bioinformatics Workflow**”) (87).

### Differential Expression and Accessibility Region Analyses Across EC Subtypes and EndMT Perturbation-Subtype Combinations

Differential expression between clusters was computed by constructing a logistic regression (LR) model predicting group membership based on the expression of a given gene in the set of cells being compared. The LR model included pro-EndMT perturbation as a latent variable and was compared to a null model using a likelihood ratio test (LRT). This was performed using Seurat FindMarkers, with “test.use = LR” and “latent.vars” set to perturbation. Differential expression between perturbation and control for each cluster was performed using pseudobulk method with DESeq2 (88). Raw RNA counts were extracted for each EndMT perturbation-subtype combination and counts, and metadata were aggregated to the sample level.

Differential accessibility between EC subtypes was performed using FindMarkers, with “test.use = LR” and latent.vars set to both the number of reads in peaks and perturbation. Finally, differential accessibility between perturbation and control for each cluster was performed using FindMarkers, with “test.use = LR” and latent.vars set to the number of reads in peaks. Bonferroni-adjusted p-values were used to determine significance at adjusted p-value < 0.05 for differential expression, and p-value < 0.005 for differential accessibility (65).

### Pathway Enrichment Analysis

Pathway enrichment analysis (PEA) was performed using Metascape (69). Top DEGs for each EC subtype or subtype-perturbation were sorted based on ascending p-value. Genes listed for each pathway were pulled from the Metacape results file, “_FINAL_GO.csv”. For heatmaps produced by metascape, top 20 or 100 pathways were pulled from Metascape .png files, “HeatmapSelectedGO.png”, “HeatmapSelectedGOParent.png”, or “HeatmapSelectedGOTop100.png”.

### Motif Enrichment Analysis

A hypergeometric test was used to test for overrepresentation of each DNA motif in the set of differentially accessible peaks compared to a background set of peaks. We tested motifs present in the Jaspar database (2020 release) (49) by first identifying which peaks contained each motif using motifmatchr R package (https://bioconductor.org/packages/motifmatchr). We computed the GC content (percentage of G and C nucleotides) for each differentially accessible peak and sampled a background set of 40,000 peaks matched for GC content (48). Per-cell motif activity scores were computed by running chromVAR (89), and visualized using Seurat (44) function FeaturePlot.

### Human Atherosclerosis scRNA-seq Public Data Download, Mapping, and Integration Across Samples

Count matrices of 17 samples taken from four different published scRNA-seq datasets were downloaded from the NCBI Gene Expression Omnibus (accessions listed in **Table S11** in the **Data Supplement**), processed using Cell Ranger (10x Genomics Cell Ranger 6.0.0) (90) with reference GRCh38 (version refdata-gex-GRCh38-2020-A, 10X Genomics), and analyzed using Seurat version 4.3.0 (44). Seurat objects were created from each dataset, and cells with < 500 counts and > 20% mitochondrial gene percent expression were excluded. Additionally, doublets were removed using DoubletFinder (91), which predicts doublets according to each real cell’s proximity in gene expression space to artificial doublets created by averaging the transcriptional profile of randomly chosen cell pairs. Next, normalization and variance stabilization, followed by PC analysis for 30 PCs were performed in Seurat (44) using default parameters. Batch effects across the 17 samples were corrected using Seurat functions (44) FindIntegrationAnchors using 10,000 anchors, followed by IntegrateData. During the integration step, cell cycle regression was performed by assigning cell cycle scores with Seurat (44) function CellCycleScoring. The *ex vivo* dataset was first visualized, and canonical markers were identified for annotating cell types using FindAllMarkers.

### Module Scoring

FindAllMarkers was used to identify top DEGs between each *ex vivo* cell subtype. Cells from the *in vitro* dataset were assigned an *ex vivo* cell subtype module score using Seurat (44) function AddModuleScore. The difference in module score between each *in vitro* EC subtype was established using Wilcoxon rank sum test with continuity correction and a two-sided alternative hypothesis.

### Comparison of *Ex Vivo* snRNA-seq Data to *In Vitro* snRNA-seq Data

Meta-analyzed *ex vivo* human scRNA-seq data and *in vitro* snRNA-seq data were compared. Gene expression values for each *ex vivo* cell subtype and *in vitro* EC subtype-perturbation were produced using the AverageExpression function in Seurat (44) (which exponentiates log data, therefore output is depth normalized in non-log space). Figure 6A was generated using hclust function in R (92). Spearman correlation was used as the distance metric. Sample clustering was performed using all significant genes (adjusted p-value < 0.05) induced and attenuated across all *in vitro* EC subtypes for each pro-EndMT perturbation versus its respective control. **Figure S8A** was made using average expression data for marker genes for each *ex vivo* cell subtype. Hierarchical clustering across *ex vivo* cell subtypes was performed using hclust function in R (92), using average expression as the distance metric for a given gene.

### GWAS SNP Enrichment Analysis

The SNPs associated with CAD were extracted from the most recent available meta-analysis (6). We utilized a matched background of SNPs pulled from 1000 Genomes Project reference panel (phase 3) (80) which were filtered using PLINK (93) v1.90b5.3 with the following settings: “--maf 0.01”, “--geno 0.05”. Mapping between genomic coordinates was performed using liftOver (81). To evaluate for enrichment in CAD-associated SNPs for each EC subtype and perturbation-subtype peak set, traseR package in R (traseR) (94) was used with the following: ‘test.method’ = “fisher”, ‘alternative’ = “greater”.

### Peak-To-Gene Linkage

We estimated a linkage score for each peak-gene pair using the LinksPeaks function in Signac (48). For each gene, we computed the Pearson correlation coefficient *r* between the gene expression and the accessibility of each peak within 500 kb of the gene TSS. For each peak, we then computed a background set of expected correlation coefficients given properties of the peak by randomly sampling 200 peaks located on a different chromosome to the gene, matched for GC content, accessibility, and sequence length (MatchRegionStats function in Signac). We then computed the Pearson correlation between the expression of the gene and the set of background peaks. A z score was computed for each peak as *z* = (*r* − *μ*)/*σ*, where *μ* was the background mean correlation coefficient and *σ* was the s.d. of the background correlation coefficients for the peak. We computed a P value for each peak using a one-sided z-test and retained peak-gene links with a p-value < 0.05 and a Pearson correlation coefficient. The results were restricted to peak regions which overlapped with significant CAD-associated SNPs (**Methods**, **“GWAS SNP Enrichment Analysis”**).

### Data Visualization

Data visualizations were performed using Seurat functions DimPlot, DotPlot, FeaturePlot, and VlnPlot. Other data visualizations were performed using ggplot2 (for stacked bar graphs) (95), UpSetR (for UpSet plots) (96), pheatmap (for DEG and DAR analysis heatmaps) and heatmap.2 (for Spearman’s rank correlation coefficient heatmap and **Figure S8A**) (97).

## DATA AVAILABILITY

Data produced in this study is made public in the GEO accession GSE228428.

## ACKNOWLEDGEMENTS

Funding for this study was provided by grants from the National Insitututes of Health through R01HL147187 (CER), R35GM137896 (DAC), F30HL162469 (MLA), T32HL7249-45 (MLA), and from the Geneen Charitable Trust Awards Program for Coronary Heart Disease Research (CER).

## REFERENCES

1. Brown JC, Gerhardt TE, Kwon E. Risk factors for coronary artery disease. StatPearls [Internet]. 2020.

2. Hajra L, Evans AI, Chen M, Hyduk SJ, Collins T, Cybulsky MI. The NF-κB signal transduction pathway in aortic endothelial cells is primed for activation in regions predisposed to atherosclerotic lesion formation. Proceedings of the National Academy of Sciences. 2000;97(16):9052–7.

3. Birdsey GM, Shah AV, Dufton N, Reynolds LE, Almagro LO, Yang Y, et al. The endothelial transcription factor ERG promotes vascular stability and growth through Wnt/β-catenin signaling. Developmental cell. 2015;32(1):82–96.

4. Marenberg ME, Risch N, Berkman LF, Floderus B, de Faire U. Genetic susceptibility to death from coronary heart disease in a study of twins. New England Journal of Medicine. 1994;330(15):1041–6.

5. Aragam KG, Jiang T, Goel A, Kanoni S, Wolford BN, Atri DS, et al. Discovery and systematic characterization of risk variants and genes for coronary artery disease in over a million participants. Nature Genetics. 2022;54(12):1803–15.

6. Tcheandjieu C, Zhu X, Hilliard AT, Clarke SL, Napolioni V, Ma S, et al. Large-scale genome-wide association study of coronary artery disease in genetically diverse populations. Nature medicine. 2022;28(8):1679–92.

7. Kessler T, Schunkert H. Coronary artery disease genetics enlightened by genome-wide association studies. Basic to Translational Science. 2021;6(7):610–23.

8. Zhao Q, Eichten A, Parveen A, Adler C, Huang Y, Wang W, et al. Single-cell transcriptome analyses reveal endothelial cell heterogeneity in tumors and changes following antiangiogenic treatment. Cancer research. 2018;78(9):2370–82.

9. Li Z, Solomonidis EG, Meloni M, Taylor RS, Duffin R, Dobie R, et al. Single-cell transcriptome analyses reveal novel targets modulating cardiac neovascularization by resident endothelial cells following myocardial infarction. European heart journal. 2019;40(30):2507–20.

10. Kalluri AS, Vellarikkal SK, Edelman ER, Nguyen L, Subramanian A, Ellinor PT, et al. Single-cell analysis of the normal mouse aorta reveals functionally distinct endothelial cell populations. Circulation. 2019;140(2):147–63.

11. Liu Z, Ruter DL, Quigley K, Tanke NT, Jiang Y, Bautch VL. Single-cell RNA sequencing reveals endothelial cell transcriptome heterogeneity under homeostatic laminar flow. Arteriosclerosis, Thrombosis, and Vascular Biology. 2021;41(10):2575–84.

12. Kalucka J, de Rooij LP, Goveia J, Rohlenova K, Dumas SJ, Meta E, et al. Single-cell transcriptome atlas of murine endothelial cells. Cell. 2020;180(4):764–79. e20.

13. Rohlenova K, Goveia J, García-Caballero M, Subramanian A, Kalucka J, Treps L, et al. Single-cell RNA sequencing maps endothelial metabolic plasticity in pathological angiogenesis. Cell metabolism. 2020;31(4):862–77. e14.

14. Zhao G, Lu H, Chang Z, Zhao Y, Zhu T, Chang L, et al. Single-cell RNA sequencing reveals the cellular heterogeneity of aneurysmal infrarenal abdominal aorta. Cardiovascular research. 2021;117(5):1402–16.

15. Xu K, Xie S, Huang Y, Zhou T, Liu M, Zhu P, et al. Cell-type transcriptome atlas of human aortic valves reveal cell heterogeneity and endothelial to mesenchymal transition involved in calcific aortic valve disease. Arteriosclerosis, thrombosis, and vascular biology. 2020;40(12):2910–21.

16. Cheng J, Gu W, Lan T, Deng J, Ni Z, Zhang Z, et al. Single-cell RNA sequencing reveals cell type- and artery type-specific vascular remodelling in male spontaneously hypertensive rats. Cardiovascular Research. 2021;117(4):1202–16.

17. Khan S, Taverna F, Rohlenova K, Treps L, Geldhof V, de Rooij L, et al. EndoDB: a database of endothelial cell transcriptomics data. Nucleic acids research. 2019;47(D1):D736–D44.

18. Andueza A, Kumar S, Kim J, Kang D-W, Mumme HL, Perez JI, et al. Endothelial reprogramming by disturbed flow revealed by single-cell RNA and chromatin accessibility study. Cell reports. 2020;33(11):108491.

19. Tombor L, John D, Glaser S, Luxan G, Forte E, Furtado M, et al. Single cell sequencing reveals endothelial plasticity with transient mesenchymal activation after myocardial infarction. European Heart Journal. 2020;41(Supplement_2):ehaa946. 3736.

20. Evrard SM, Lecce L, Michelis KC, Nomura-Kitabayashi A, Pandey G, Purushothaman K-R, et al. Endothelial to mesenchymal transition is common in atherosclerotic lesions and is associated with plaque instability. Nature communications. 2016;7(1):1–16.

21. Chen P-Y, Qin L, Barnes C, Charisse K, Yi T, Zhang X, et al. FGF regulates TGF-β signaling and endothelial-to-mesenchymal transition via control of let-7 miRNA expression. Cell reports. 2012;2(6):1684–96.

22. Chen P-Y, Qin L, Baeyens N, Li G, Afolabi T, Budatha M, et al. Endothelial-to-mesenchymal transition drives atherosclerosis progression. The Journal of clinical investigation. 2015;125(12):4514–28.

23. Moonen J-RA, Lee ES, Schmidt M, Maleszewska M, Koerts JA, Brouwer LA, et al. Endothelial-to-mesenchymal transition contributes to fibro-proliferative vascular disease and is modulated by fluid shear stress. Cardiovascular research. 2015;108(3):377–86.

24. Yuan L, Chan GC, Beeler D, Janes L, Spokes KC, Dharaneeswaran H, et al. A role of stochastic phenotype switching in generating mosaic endothelial cell heterogeneity. Nature communications. 2016;7(1):10160.

25. Turgeon PJ, Chan GC, Chen L, Jamal AN, Yan MS, Ho J, et al. Epigenetic heterogeneity and mitotic heritability prime endothelial cell gene induction. The Journal of Immunology. 2020;204(5):1173–87.

26. Pan H, Xue C, Auerbach BJ, Fan J, Bashore AC, Cui J, et al. Single-cell genomics reveals a novel cell state during smooth muscle cell phenotypic switching and potential therapeutic targets for atherosclerosis in mouse and human. Circulation. 2020;142(21):2060–75.

27. Alsaigh T, Evans D, Frankel D, Torkamani A. Decoding the transcriptome of calcified atherosclerotic plaque at single-cell resolution. Communications biology. 2022;5(1):1–17.

28. Chowdhury RR, D’Addabbo J, Huang X, Veizades S, Sasagawa K, Louis DM, et al. Human Coronary Plaque T Cells Are Clonal and Cross-React to Virus and Self. Circulation Research. 2022;130(10):1510–30.

29. Wirka RC, Wagh D, Paik DT, Pjanic M, Nguyen T, Miller CL, et al. Atheroprotective roles of smooth muscle cell phenotypic modulation and the TCF21 disease gene as revealed by single-cell analysis. Nature medicine. 2019;25(8):1280–9.

30. van Meeteren LA, Ten Dijke P. Regulation of endothelial cell plasticity by TGF-β. Cell and tissue research. 2012;347(1):177–86.

31. Bujak M, Dobaczewski M, Chatila K, Mendoza LH, Li N, Reddy A, et al. Interleukin-1 receptor type I signaling critically regulates infarct healing and cardiac remodeling. The American journal of pathology. 2008;173(1):57–67.

32. Bujak M, Frangogiannis NG. The role of IL-1 in the pathogenesis of heart disease. Archivum immunologiae et therapiae experimentalis. 2009;57(3):165–76.

33. Maleszewska M, Moonen J-RA, Huijkman N, van de Sluis B, Krenning G, Harmsen MC. IL-1β and TGFβ2 synergistically induce endothelial to mesenchymal transition in an NFκB-dependent manner. Immunobiology. 2013;218(4):443–54.

34. Chaudhuri V, Zhou L, Karasek M. Inflammatory cytokines induce the transformation of human dermal microvascular endothelial cells into myofibroblasts: a potential role in skin fibrogenesis. Journal of cutaneous pathology. 2007;34(2):146–53.

35. Sánchez-Duffhues G, García de Vinuesa A, van de Pol V, Geerts ME, de Vries MR, Janson SG, et al. Inflammation induces endothelial-to-mesenchymal transition and promotes vascular calcification through downregulation of BMPR2. The Journal of pathology. 2019;247(3):333–46.

36. Ridker PM, Everett BM, Thuren T, MacFadyen JG, Chang WH, Ballantyne C, et al. Antiinflammatory therapy with canakinumab for atherosclerotic disease. New England journal of medicine. 2017;377(12):1119–31.

37. Sperone A, Dryden NH, Birdsey GM, Madden L, Johns M, Evans PC, et al. The transcription factor Erg inhibits vascular inflammation by repressing NF-κB activation and proinflammatory gene expression in endothelial cells. Arteriosclerosis, thrombosis, and vascular biology. 2011;31(1):142–50.

38. Fish JE, Cantu Gutierrez M, Dang LT, Khyzha N, Chen Z, Veitch S, et al. Dynamic regulation of VEGF-inducible genes by an ERK/ERG/p300 transcriptional network. Development. 2017;144(13):2428–44.

39. Lathen C, Zhang Y, Chow J, Singh M, Lin G, Nigam V, et al. ERG-APLNR axis controls pulmonary venule endothelial proliferation in pulmonary veno-occlusive disease. Circulation. 2014;130(14):1179–91.

40. Vijayaraj P, Le Bras A, Mitchell N, Kondo M, Juliao S, Wasserman M, et al. Erg is a crucial regulator of endocardial-mesenchymal transformation during cardiac valve morphogenesis. Development. 2012;139(21):3973–85.

41. Hogan NT, Whalen MB, Stolze LK, Hadeli NK, Lam MT, Springstead JR, et al. Transcriptional networks specifying homeostatic and inflammatory programs of gene expression in human aortic endothelial cells. Elife. 2017;6:e22536.

42. Navab M, Hough GP, Stevenson LW, Drinkwater DC, Laks H, Fogelman AM. Monocyte migration into the subendothelial space of a coculture of adult human aortic endothelial and smooth muscle cells. The Journal of clinical investigation. 1988;82(6):1853–63.

43. Genomics x. Chromium Next GEM Single Cell Multiome ATAC + Gene Expression. Revision F edAugust 2022.

44. Hao Y, Hao S, Andersen-Nissen E, Mauck III WM, Zheng S, Butler A, et al. Integrated analysis of multimodal single-cell data. Cell. 2021;184(13):3573–87. e29.

45. Dahal S, Huang P, Murray BT, Mahler GJ. Endothelial to mesenchymal transformation is induced by altered extracellular matrix in aortic valve endothelial cells. Journal of Biomedical Materials Research Part A. 2017;105(10):2729–41.

46. Kovacic JC, Dimmeler S, Harvey RP, Finkel T, Aikawa E, Krenning G, et al. Endothelial to mesenchymal transition in cardiovascular disease: JACC state-of-the-art review. Journal of the American College of Cardiology. 2019;73(2):190–209.

47. Bondareva O, Rodríguez-Aguilera JR, Oliveira F, Liao L, Rose A, Gupta A, et al. Single-cell profiling of vascular endothelial cells reveals progressive organ-specific vulnerabilities during obesity. Nature Metabolism. 2022:1–20.

48. Stuart T, Srivastava A, Madad S, Lareau CA, Satija R. Single-cell chromatin state analysis with Signac. Nature methods. 2021;18(11):1333–41.

49. Fornes O, Castro-Mondragon JA, Khan A, Van der Lee R, Zhang X, Richmond PA, et al. JASPAR 2020: update of the open-access database of transcription factor binding profiles. Nucleic acids research. 2020;48(D1):D87–D92.

50. Nagai N, Ohguchi H, Nakaki R, Matsumura Y, Kanki Y, Sakai J, et al. Downregulation of ERG and FLI1 expression in endothelial cells triggers endothelial-to-mesenchymal transition. PLoS genetics. 2018;14(11):e1007826.

51. Örd T, Õunap K, Stolze LK, Aherrahrou R, Nurminen V, Toropainen A, et al. Single-cell epigenomics and functional fine-mapping of atherosclerosis GWAS loci. Circulation research. 2021;129(2):240–58.

52. Zhang L, Tang C, Zhang M, Tong X, Xie Y, Yan R, et al. Single cell meta-analysis of EndMT and EMT state in COVID-19. Frontiers in immunology. 2022;13.

53. Deng G, Zhang L, Wang C, Wang S, Xu J, Dong J, et al. AGEs-RAGE axis causes endothelial-to-mesenchymal transition in early calcific aortic valve disease via TGF-β1 and BMPR2 signaling. Experimental Gerontology. 2020;141:111088.

54. Ekiz HA, Conley CJ, Stephens WZ, O’Connell RM. CIPR: a web-based R/shiny app and R package to annotate cell clusters in single cell RNA sequencing experiments. BMC bioinformatics. 2020;21(1):1–15.

55. Ricciotti E, FitzGerald GA. Prostaglandins and inflammation. Arteriosclerosis, thrombosis, and vascular biology. 2011;31(5):986–1000.

56. Levi M, van der Poll T, Büller HR. Bidirectional relation between inflammation and coagulation. Circulation. 2004;109(22):2698–704.

57. Ni C, Buszczak M, editors. The homeostatic regulation of ribosome biogenesis. Seminars in Cell & Developmental Biology; 2022: Elsevier.

58. Suárez Y, Fernández-Hernando C, Pober JS, Sessa WC. Dicer dependent microRNAs regulate gene expression and functions in human endothelial cells. Circulation research. 2007;100(8):1164–73.

59. Krizbai IA, Gasparics Á, Nagyőszi P, Fazakas C, Molnár J, Wilhelm I, et al. Endothelial-mesenchymal transition of brain endothelial cells: possible role during metastatic extravasation. PloS one. 2015;10(3):e0119655.

60. Zhao G, Lu H, Liu Y, Zhao Y, Zhu T, Garcia-Barrio MT, et al. Single-cell transcriptomics reveals endothelial plasticity during diabetic atherogenesis. Frontiers in cell and developmental biology. 2021:1213.

61. Pinto MT, Melo FUF, Malta TM, Rodrigues ES, Placa JR, Silva Jr WA, et al. Endothelial cells from different anatomical origin have distinct responses during SNAIL/TGF-β2-mediated endothelial-mesenchymal transition. American journal of translational research. 2018;10(12):4065.

62. Stenmark KR, Frid M, Perros F. Endothelial-to-mesenchymal transition: an evolving paradigm and a promising therapeutic target in PAH. Am Heart Assoc; 2016. p. 1734–7.

63. Gole S, Tkachenko S, Masannat T, Baylis RA, Cherepanova OA. Endothelial-to-Mesenchymal Transition in Atherosclerosis: Friend or Foe? Cells. 2022;11(19):2946.

64. Lee Y-H, Albig AR, Regner M, Schiemann BJ, Schiemann WP. Fibulin-5 initiates epithelial– mesenchymal transition (EMT) and enhances EMT induced by TGF-β in mammary epithelial cells via a MMP-dependent mechanism. Carcinogenesis. 2008;29(12):2243–51.

65. Benjamini Y, Hochberg Y. Controlling the false discovery rate: a practical and powerful approach to multiple testing. Journal of the Royal statistical society: series B (Methodological). 1995;57(1):289–300.

66. Drobni ZD, Kolossvary M, Karady J, Jermendy AL, Tarnoki AD, Tarnoki DL, et al. Heritability of coronary artery disease: Insights from a classical twin study. Circulation: Cardiovascular Imaging. 2022;15(3):e013348.

67. McPherson R, Tybjaerg-Hansen A. Genetics of coronary artery disease. Circulation research. 2016;118(4):564–78.

68. Cao J, Cusanovich DA, Ramani V, Aghamirzaie D, Pliner HA, Hill AJ, et al. Joint profiling of chromatin accessibility and gene expression in thousands of single cells. Science. 2018;361(6409):1380-5.

69. Zhou Y, Zhou B, Pache L, Chang M, Khodabakhshi AH, Tanaseichuk O, et al. Metascape provides a biologist-oriented resource for the analysis of systems-level datasets. Nature communications. 2019;10(1):1–10.

70. Liu T, Zou X-Z, Huang N, Ge X-Y, Yao M-Z, Liu H, et al. miR-27a promotes endothelial-mesenchymal transition in hypoxia-induced pulmonary arterial hypertension by suppressing BMP signaling. Life sciences. 2019;227:64–73.

71. Yang W, Ng FL, Chan K, Pu X, Poston RN, Ren M, et al. Coronary-heart-disease-associated genetic variant at the COL4A1/COL4A2 locus affects COL4A1/COL4A2 expression, vascular cell survival, atherosclerotic plaque stability and risk of myocardial infarction. PLoS genetics. 2016;12(7):e1006127.

72. Woodfin A, Voisin M-B, Nourshargh S. PECAM-1: a multi-functional molecule in inflammation and vascular biology. Arteriosclerosis, thrombosis, and vascular biology. 2007;27(12):2514–23.

73. Rodor J, Chen SH, Scanlon JP, Monteiro JP, Caudrillier A, Sweta S, et al. Single-cell RNA sequencing profiling of mouse endothelial cells in response to pulmonary arterial hypertension. Cardiovascular Research. 2022;118(11):2519–34.

74. Medici D, Potenta S, Kalluri R. Transforming growth factor-β2 promotes Snail-mediated endothelial–mesenchymal transition through convergence of Smad-dependent and Smad-independent signalling. Biochemical Journal. 2011;437(3):515–20.

75. Stolze LK, Conklin AC, Whalen MB, Rodríguez ML, Õunap K, Selvarajan I, et al. Systems Genetics in Human Endothelial Cells Identifies Non-coding Variants Modifying Enhancers, Expression, and Complex Disease Traits. The American Journal of Human Genetics. 2020;106(6):748–63.

76. Toropainen A, Stolze LK, Örd T, Whalen MB, Torrell PM, Link VM, et al. Functional noncoding SNPs in human endothelial cells fine-map vascular trait associations. Genome Research. 2022;32(3):409–24.

77. Schneider CA, Rasband WS, Eliceiri KW. NIH Image to ImageJ: 25 years of image analysis. Nature methods. 2012;9(7):671–5.

78. Genomics x. Nuclei Isolation for Single Cell Multiome ATAC + Gene Expression Sequencing. Revision C ed 2022.

79. Howie BN, Donnelly P, Marchini J. A flexible and accurate genotype imputation method for the next generation of genome-wide association studies. PLoS genetics. 2009;5(6):e1000529.

80. Consortium GP, Auton A, Brooks L, Durbin R, Garrison E, Kang H. A global reference for human genetic variation. Nature. 2015;526(7571):68-74.

81. Kuhn RM, Haussler D, Kent WJ. The UCSC genome browser and associated tools. Briefings in bioinformatics. 2013;14(2):144–61.

82. Danecek P, Auton A, Abecasis G, Albers CA, Banks E, DePristo MA, et al. The variant call format and VCFtools. Bioinformatics. 2011;27(15):2156–8.

83. Danecek P, Bonfield JK, Liddle J, Marshall J, Ohan V, Pollard MO, et al. Twelve years of SAMtools and BCFtools. Gigascience. 2021;10(2):giab008.

84. Kang HM, Subramaniam M, Targ S, Nguyen M, Maliskova L, McCarthy E, et al. Multiplexed droplet single-cell RNA-sequencing using natural genetic variation. Nature biotechnology. 2018;36(1):89–94.

85. Luecken MD, Theis FJ. Current best practices in single-cell RNA-seq analysis: a tutorial. Molecular systems biology. 2019;15(6):e8746.

86. Heidecker B, Lamirault G, Kasper EK, Wittstein IS, Champion HC, Breton E, et al. The gene expression profile of patients with new-onset heart failure reveals important gender-specific differences. European heart journal. 2010;31(10):1188–96.

87. Yan F, Powell DR, Curtis DJ, Wong NC. From reads to insight: a hitchhiker’s guide to ATAC-seq data analysis. Genome biology. 2020;21:1–16.

88. Love MI, Huber W, Anders S. Moderated estimation of fold change and dispersion for RNA-seq data with DESeq2. Genome biology. 2014;15(12):1–21.

89. Schep AN, Wu B, Buenrostro JD, Greenleaf WJ. chromVAR: inferring transcription-factor-associated accessibility from single-cell epigenomic data. Nature methods. 2017;14(10):975–8.

90. Zheng GX, Terry JM, Belgrader P, Ryvkin P, Bent ZW, Wilson R, et al. Massively parallel digital transcriptional profiling of single cells. Nature communications. 2017;8(1):14049.

91. McGinnis CS, Murrow LM, Gartner ZJ. DoubletFinder: doublet detection in single-cell RNA sequencing data using artificial nearest neighbors. Cell systems. 2019;8(4):329–37. e4.

92. Murtagh F, Legendre P. Ward’s hierarchical agglomerative clustering method: which algorithms implement Ward’s criterion? Journal of classification. 2014;31(3):274–95.

93. Purcell S, Neale B, Todd-Brown K, Thomas L, Ferreira MA, Bender D, et al. PLINK: a tool set for whole-genome association and population-based linkage analyses. The American journal of human genetics. 2007;81(3):559–75.

94. Chen L, Qin ZS. traseR: an R package for performing trait-associated SNP enrichment analysis in genomic intervals. Bioinformatics. 2016;32(8):1214–6.

95. Villanueva RAM, Chen ZJ. ggplot2: elegant graphics for data analysis. Taylor & Francis; 2019.

96. Conway EM, Collen D, Carmeliet P. Molecular mechanisms of blood vessel growth. Cardiovascular research. 2001;49(3):507–21.

97. Warnes GR, Bolker B, Bonebakker L, Gentleman R, Liaw WHA, Lumley T, et al. Gplots: various R programming tools for plotting data. 2016. R package version. 2014;2(0).

