## Supplementary Figures for "Single cell ‘omic profiles of human aortic endothelial cells *in vitro* and human atherosclerotic lesions *ex vivo* reveals heterogeneity of endothelial subtype and response to activating perturbations"

A

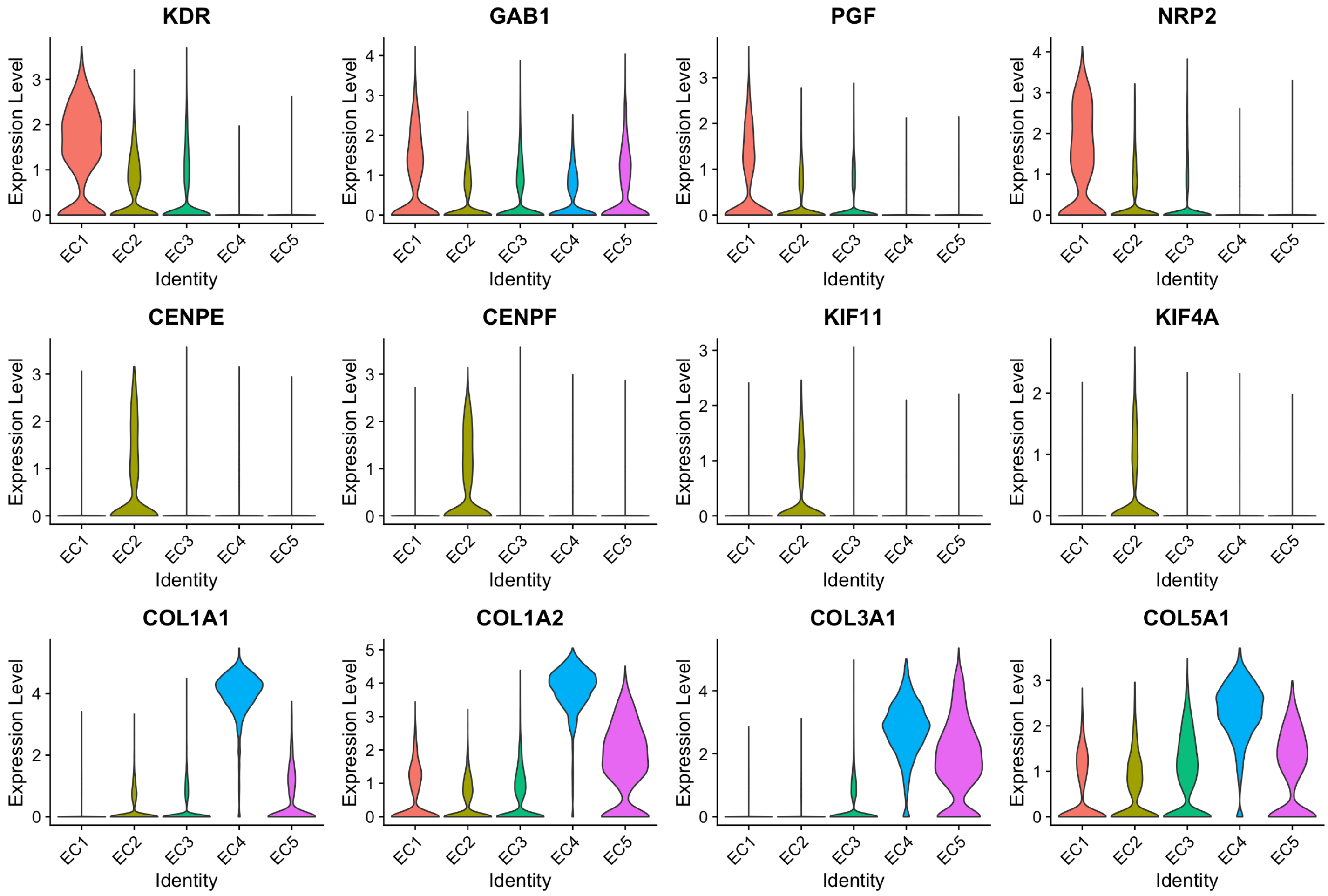

KDR
GAB1
PGF
NRP2
CENPE
CENPF
KIF11
KIF4A
COL1A1
COL1A2
COL3A1
COL5A1
Expression Level
Identity
EC1
EC2
EC3
EC4
EC5

B

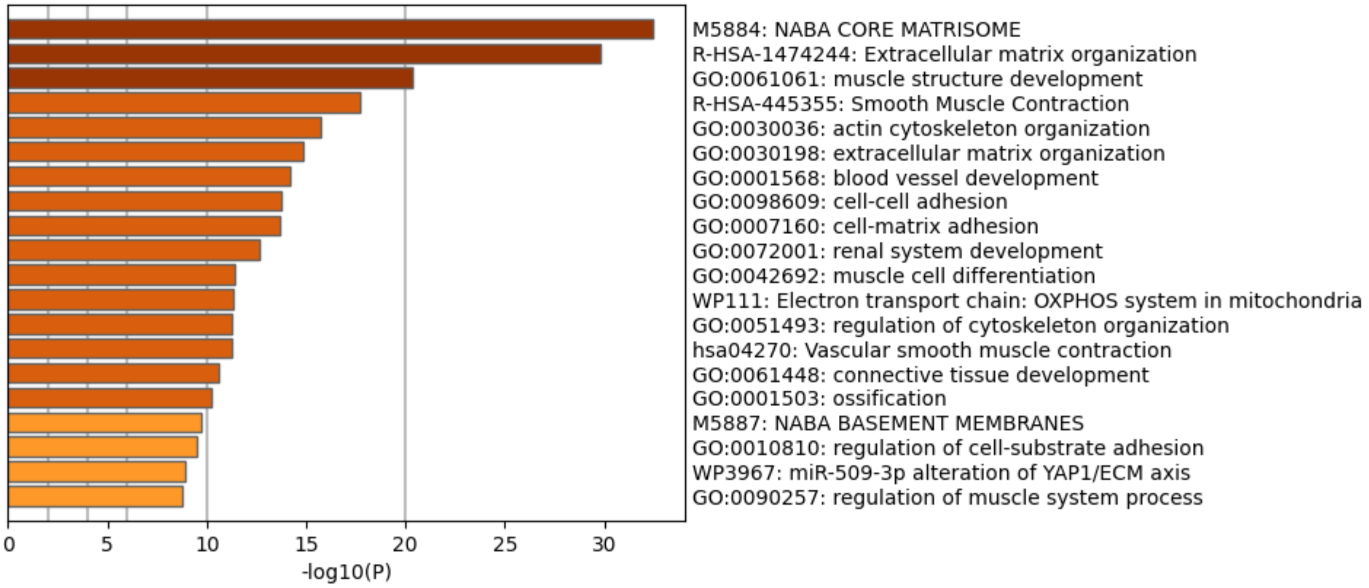


C

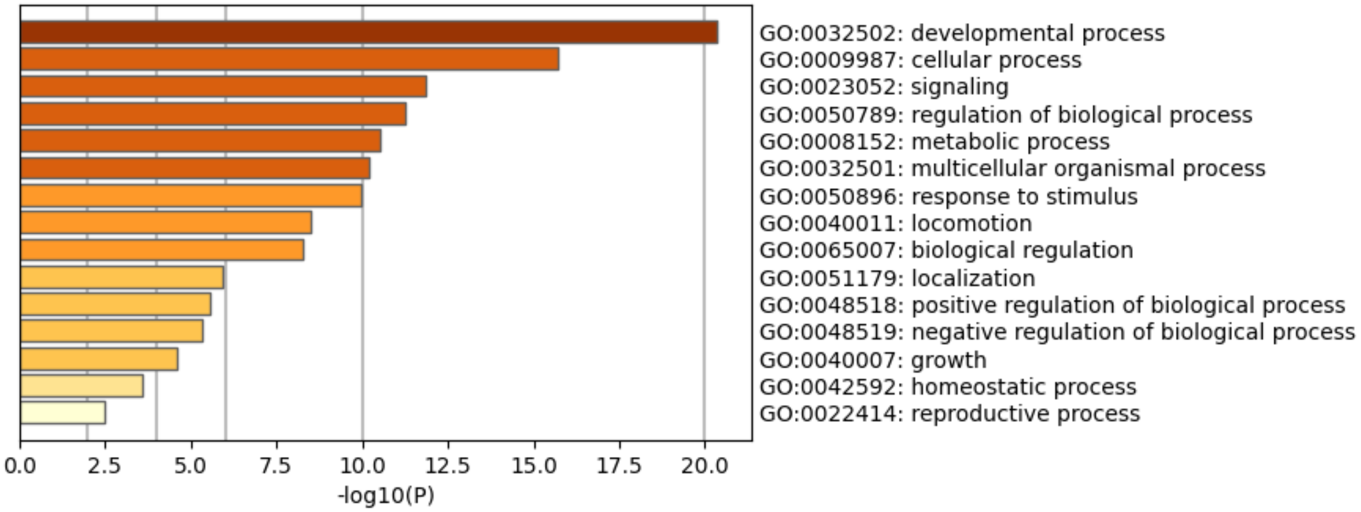


**Supplementary Figure 1 (Figure S1) – Related to Figure 1 | (A)**, Violin plots of representative marker genes for angiogenic EC1 (*KDR*, *GAB1*, *PGF*, *NRP1*), proliferative EC2 (*CENPE*, *CENPF*, *KIF11*, *KIF4A*), and mesenchymal EC4 (*COL1A1*, *COL1A2*, *COL3A1*, *COL5A1*) sub-phenotypes. **(B)**, Top 20 pathway enrichment analysis (PEA) results from submitting top 200 differentially expressed genes (DEGs; by ascending p-value) regulated in EC3 versus EC1-2 and EC4-5. **(C)**, Top 20 Gene Ontology (GO) PEA results from submitting top 200 DEGs (by ascending p-value) regulated in EC3 versus EC1-2 and EC4-5.

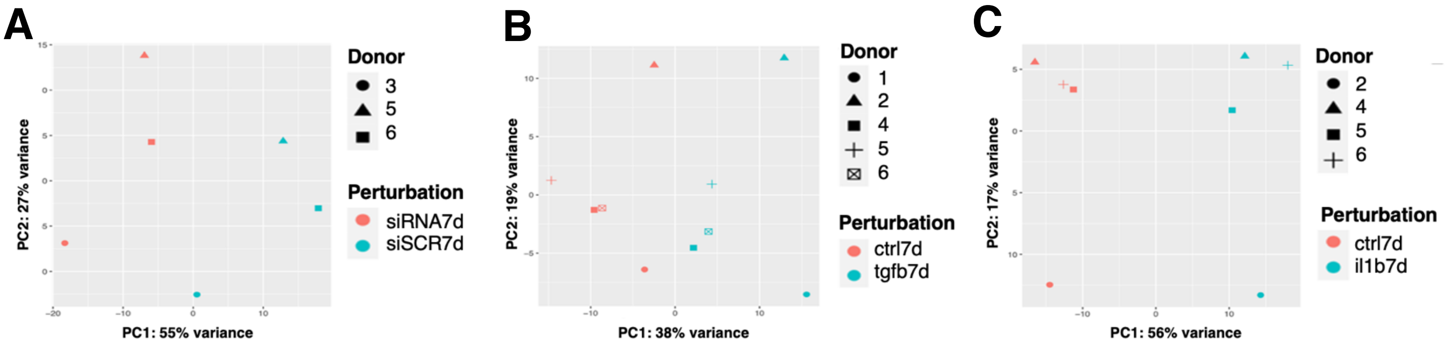


**Supplementary Figure 2 (Figure S2) – Related to Figure 3 I (A)**, Principal component analysis (PCA) of EC1-4 snRNA-seq samples +/- siERG or control across donor replicates. **(B)**, PCA of EC1-4 snRNA-seq samples +/- TGFB2 or control across donor replicates. **(C)**, PCA of EC1-4 snRNA-seq samples +/- IL1B or control across donor replicates.

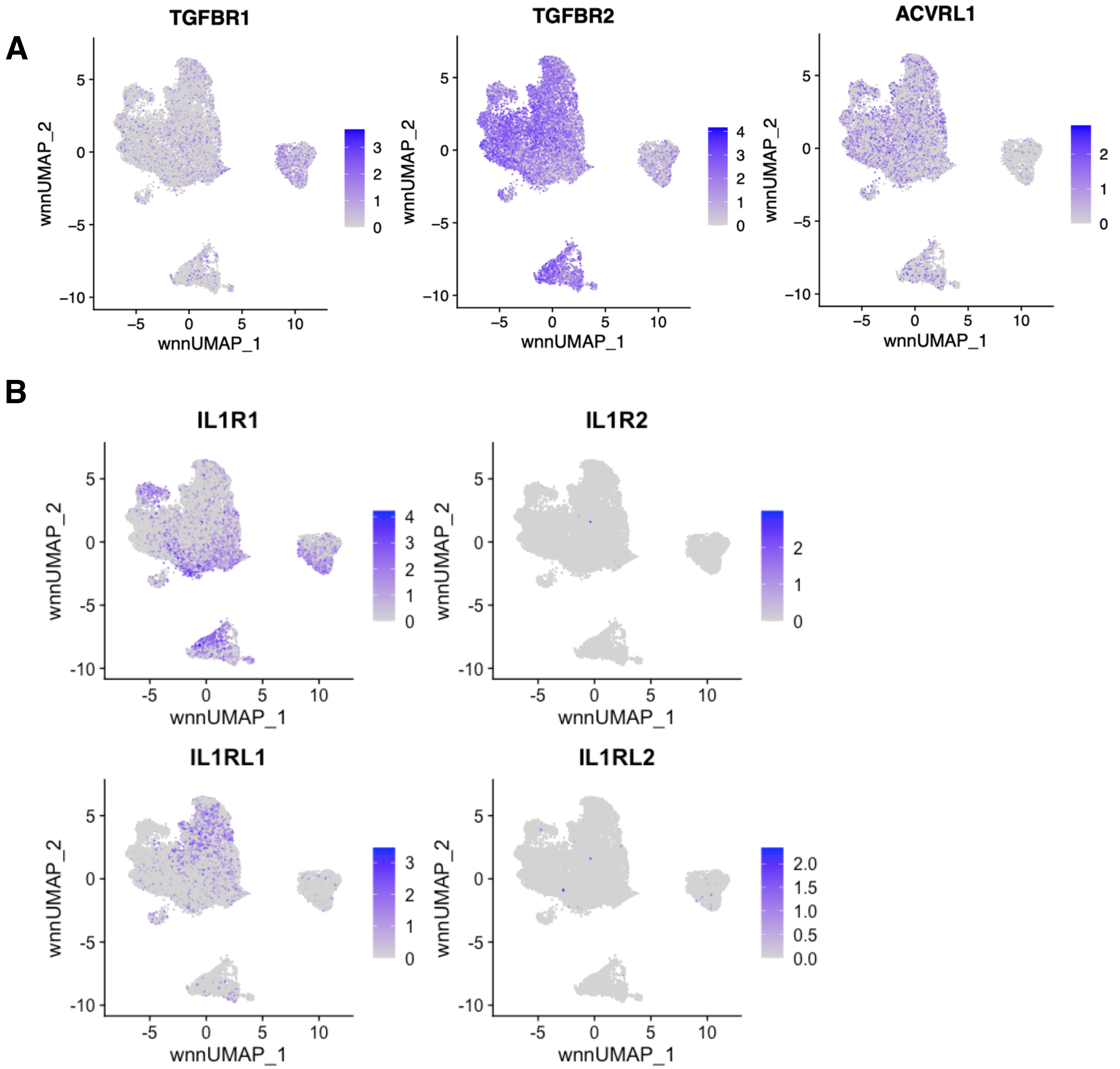


**Supplementary Figure 3 (Figure S3) – Related to Figure 3 l (A)**, Feature plots of expression of TGFB pathway receptors: *TGFBR1*, *TGFBR2*, and *ACVRL1*. **(B)**, Feature plots of IL1B pathway receptors: *IL1R1*, *IL1R2*, *IL1RL1*, and *IL1RL2*.

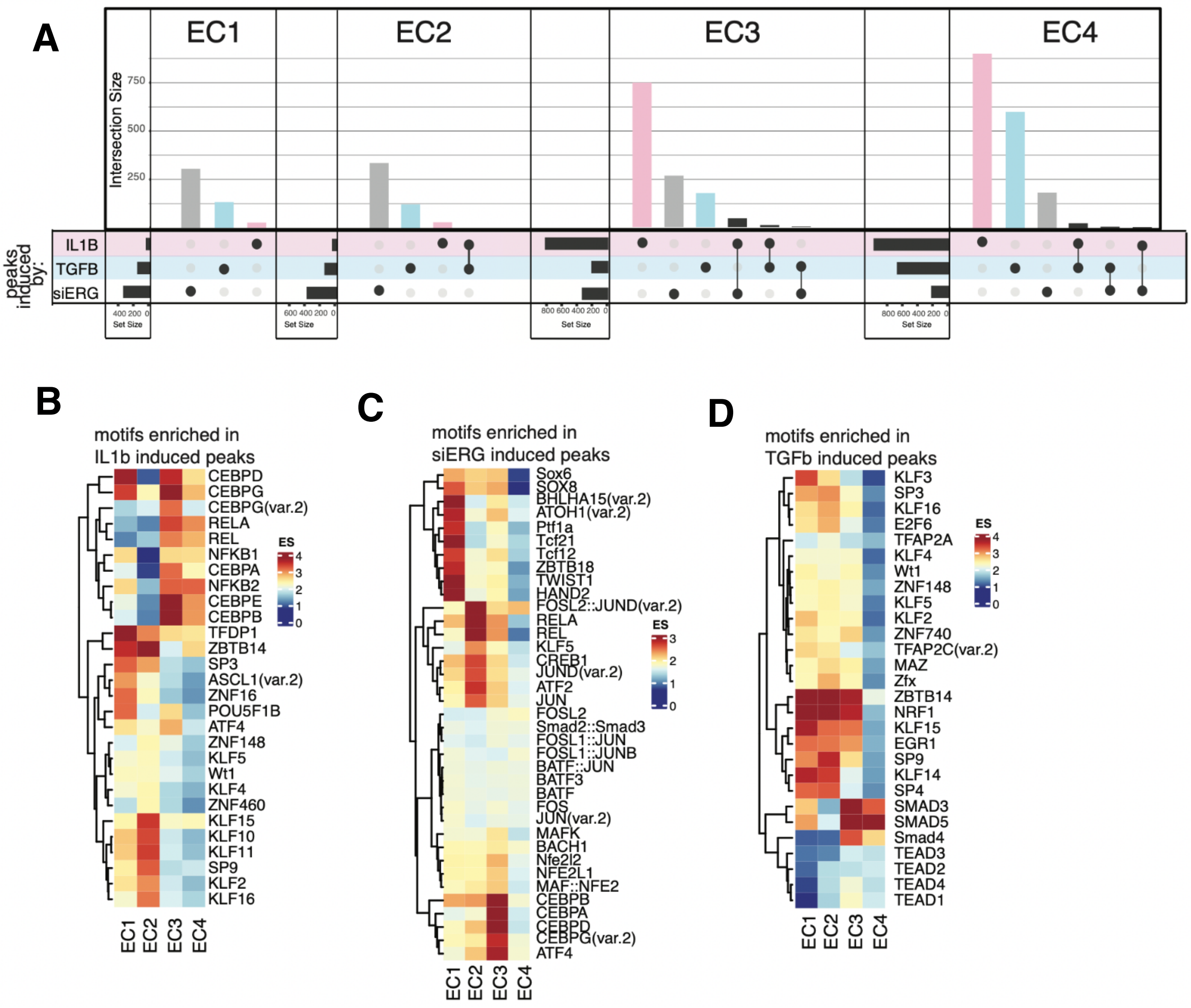


**Supplementary Figure 4 (Figure S4) | (A)**, Upset plot of induced peaks for siERG (grey), IL1B (pink), and TGFB2 (blue) across EC1, EC2, EC3, and EC4. Upset plots visualize intersections between sets in a matrix, where the columns of the matrix correspond to the sets, and the rows correspond to the intersections. Intersection size represents the number of genes at each intersection. **(B)**, Heatmap of top motifs enriched in IL1B-induced peaks. Top TFs for each EC subtype are selected based on ascending p-value. Rows (TFs) and columns (EC subtype) are clustered based on enrichment score (ES). **(C)**, Heatmap of top motifs enriched in siERG-induced peaks. **(D)**, Heatmap of top motifs enriched in TGFB2-induced peaks.

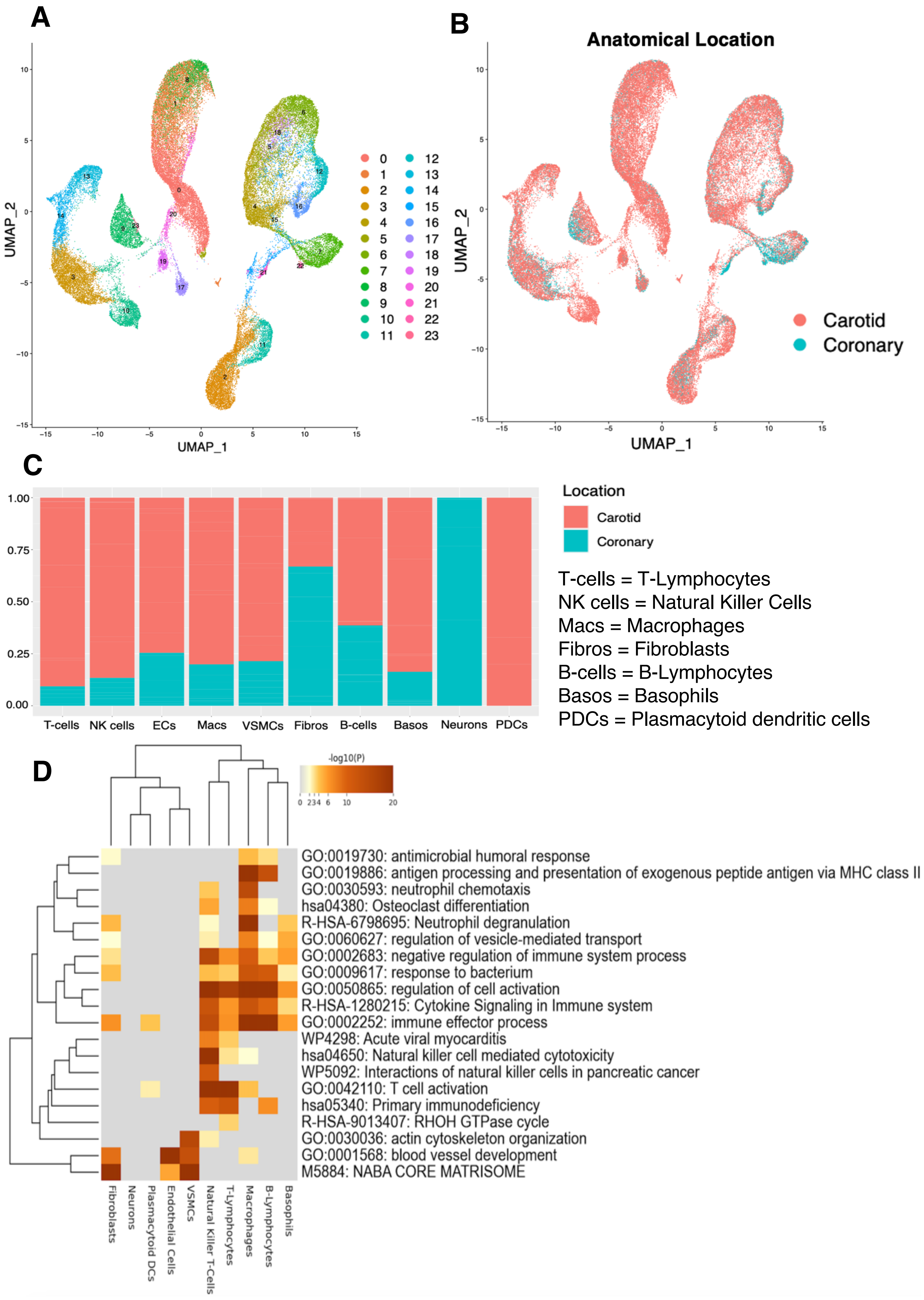

A
UMAP_2
UMAP_1
B
Anatomical Location
UMAP_2
UMAP_1
Carotid
Coronary
C
Location
Carotid
Coronary
T-cells = T-Lymphocytes
NK cells = Natural Killer Cells
Macs = Macrophages
Fibros = Fibroblasts
B-cells = B-Lymphocytes
Basos = Basophils
PDCs = Plasmacytoid dendritic cells
T-cells
NK cells
ECs
Macs
VSMCs
Fibros
B-cells
Basos
Neurons
PDCs
D
-log10(P)
GO:0019730: antimicrobial humoral response
GO:0019886: antigen processing and presentation of exogenous peptide antigen via MHC class II
GO:0030593: neutrophil chemotaxis
hsa04380: Osteoclast differentiation
R-HSA-6798695: Neutrophil degranulation
GO:0060627: regulation of vesicle-mediated transport
GO:0002683: negative regulation of immune system process
GO:0009617: response to bacterium
GO:0050865: regulation of cell activation
R-HSA-1280215: Cytokine Signaling in Immune system
GO:0002252: immune effector process
WP4298: Acute viral myocarditis
hsa04650: Natural killer cell mediated cytotoxicity
WP5092: Interactions of natural killer cells in pancreatic cancer
GO:0042110: T cell activation
hsa05340: Primary immunodeficiency
R-HSA-9013407: RHOH GTPase cycle
GO:0030036: actin cytoskeleton organization
GO:0001568: blood vessel development
M5884: NABA CORE MATRISOME
Fibroblasts
Neurons
Plasmacytoid DCs
Endothelial Cells
VSMCs
Natural Killer T-Cells
T-Lymphocytes
Macrophages
B-Lymphocytes
Basophils

E

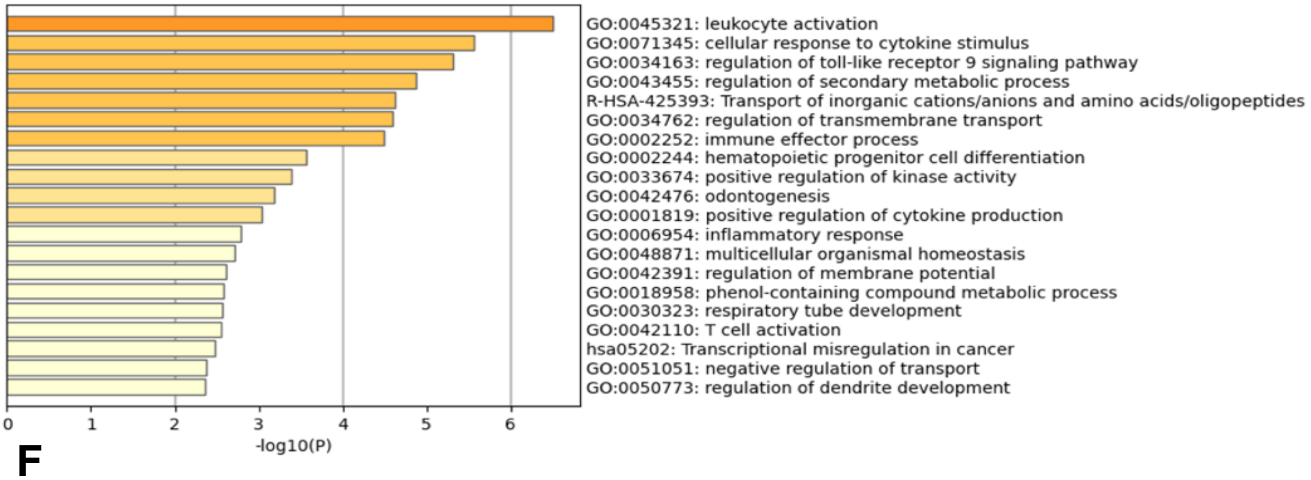


F

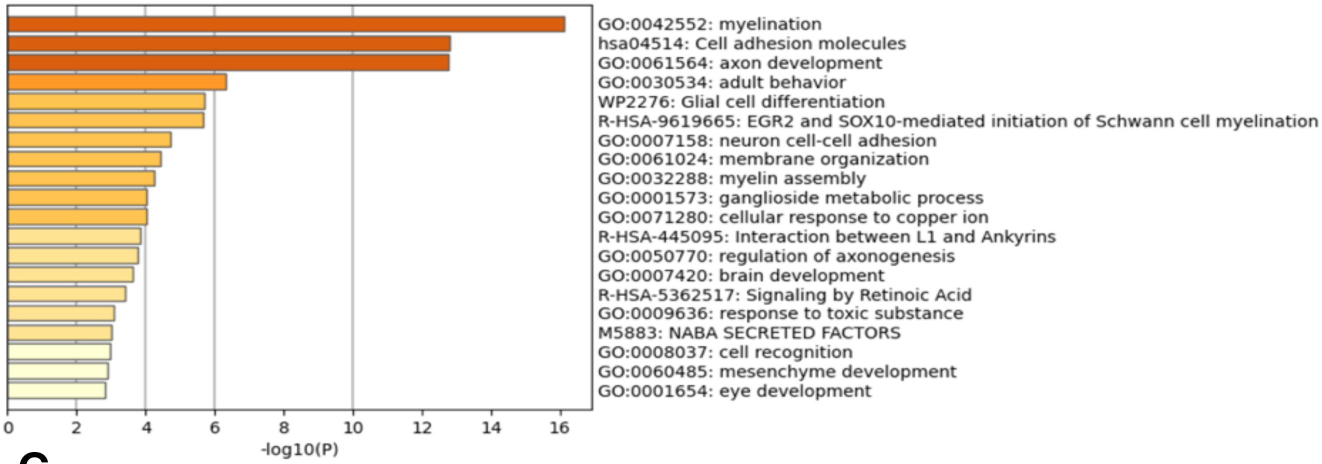


G

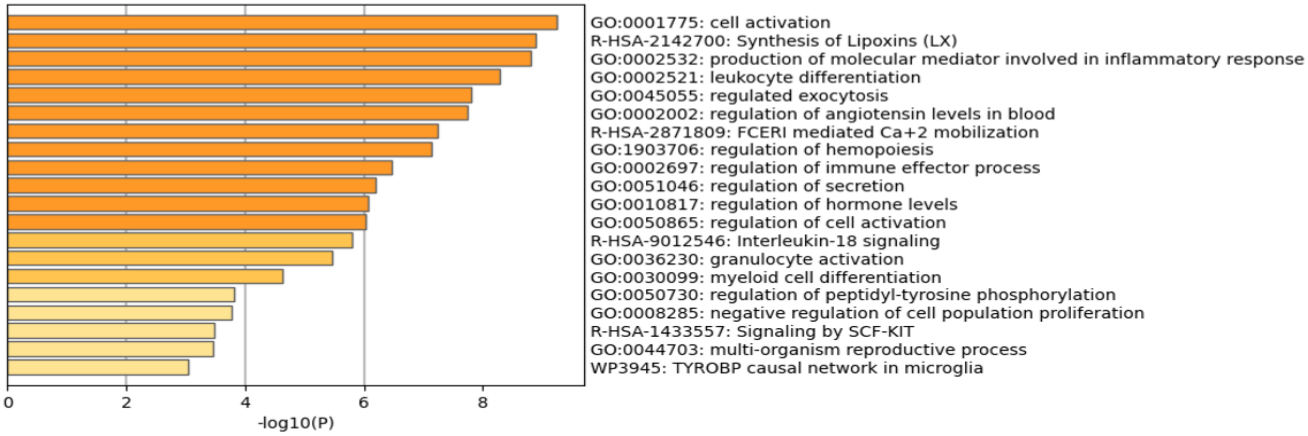


**Supplementary Figure 5 (Figure S5) – Related to Figure 5 | (A)**, UMAP displaying original clusters formed from scRNA-seq data taken from 17 samples across 4 studies of human *ex vivo* atherosclerotic plaques. Colors denote different clusters. **(B)**, UMAP from (A). Colors denote anatomical location from which cells derived. **(C)**, Stacked bar graph showing the distribution of anatomic location (red denoting carotid, blue denoting coronary arteries) from which cells derived. **(D)**, Heatmap of PEA results from submitting top 100 DEGs (by ascending p-value) between *ex vivo* cell types. Rows (pathways) and columns (cell subtypes) are clustered based on $-\text{Log}_{10}(\text{P})$**. (E)**, PEA of the top 100 DEGs (by ascending p-value) for PDCs. **(F)**, PEA of the top 100 DEGs (by ascending p-value) for neurons. **(G)**, PEA of the top 100 DEGs (by ascending p-value) for basophils. Adjusted p-value < 0.05 for DEGs submitted in D-G.

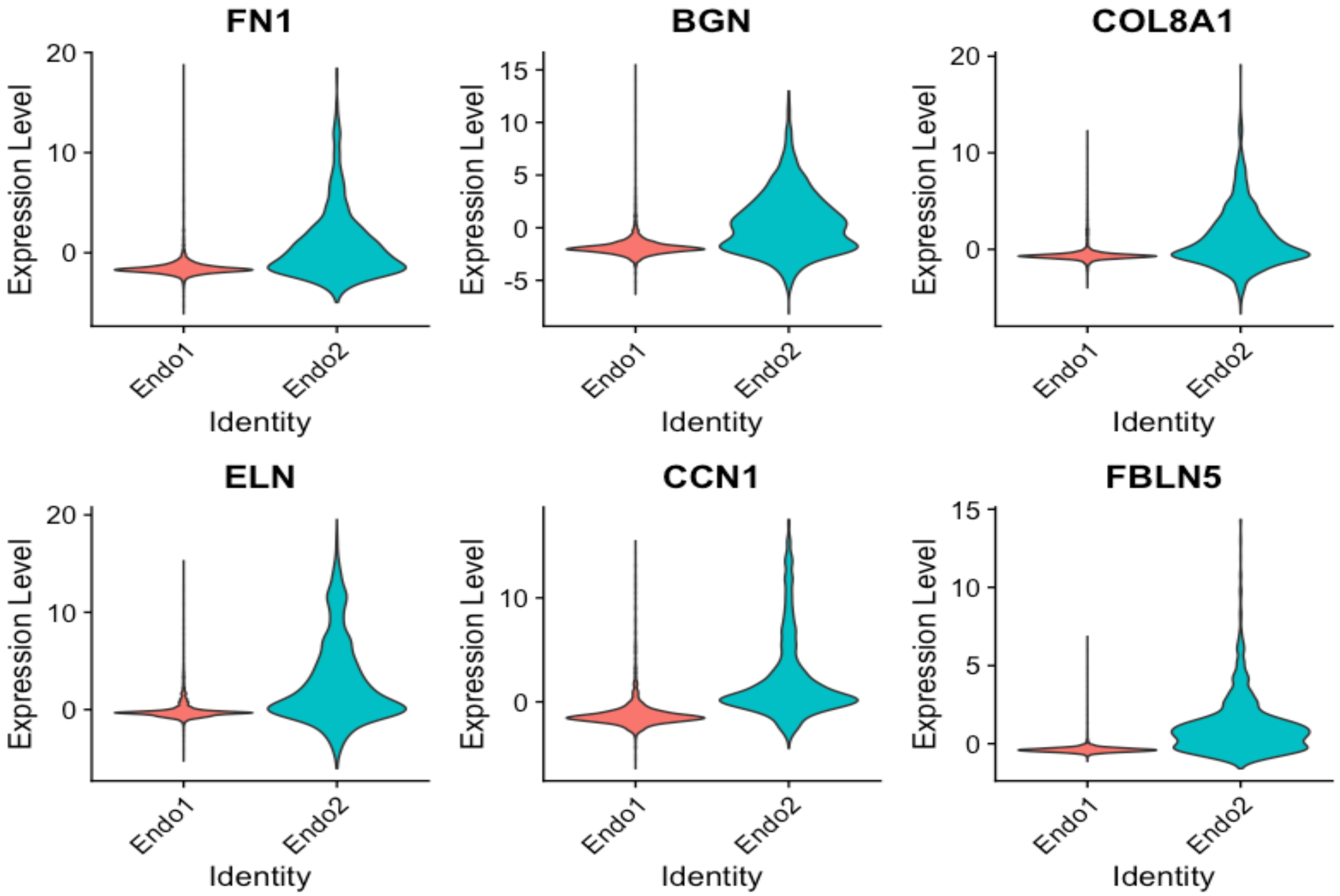


**Supplementary Figure 6 (Figure S6) – Related to Figure 5 l** Violin plots displaying upregulation of several EndMT markers in Endo2, compared to Endo1, including: *FN1*, *BGN*, *COL8A1*, *ELN*, *CCN1*, *FBLN5*.

A

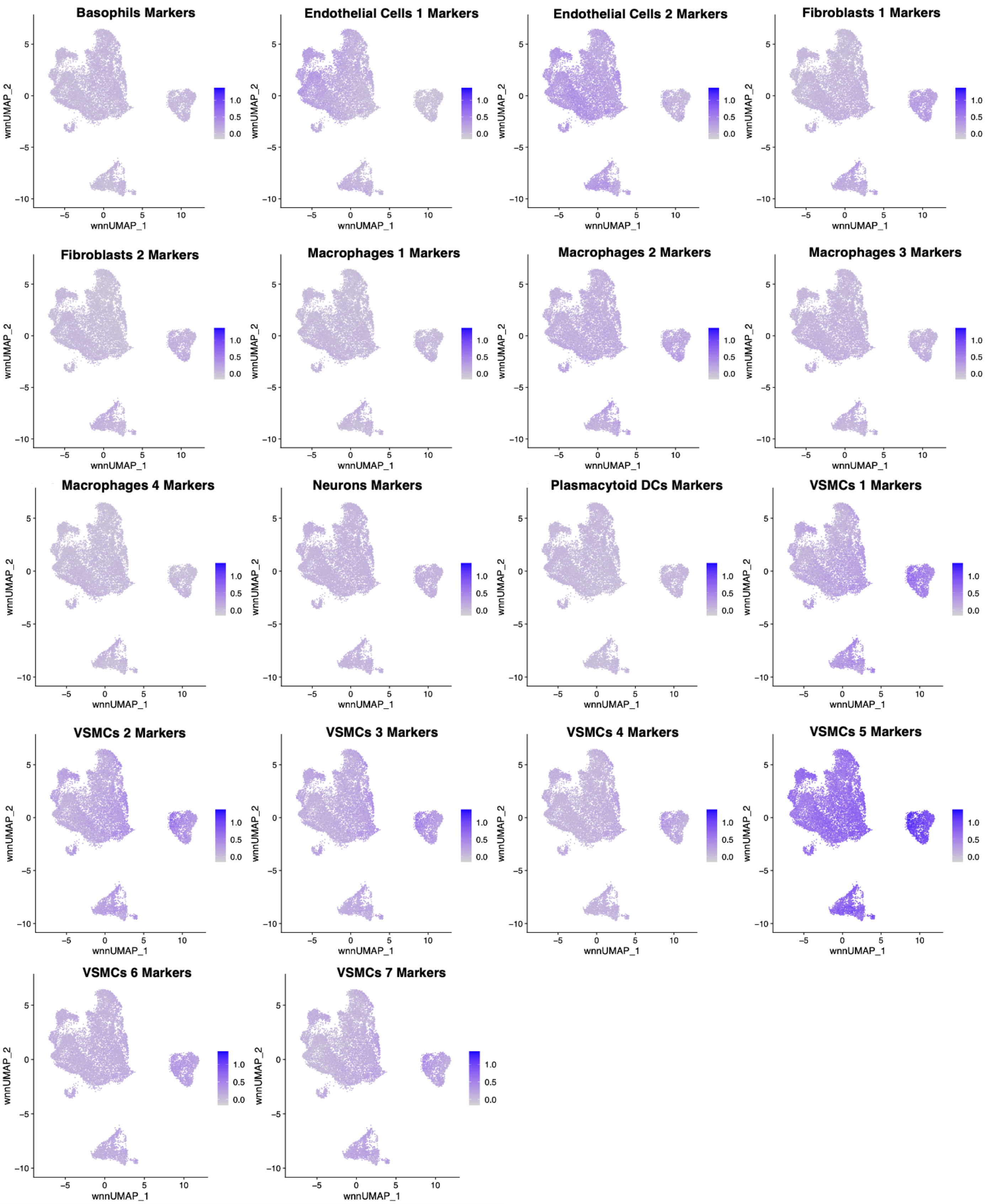

Basophils Markers
Endothelial Cells 1 Markers
Endothelial Cells 2 Markers
Fibroblasts 1 Markers
Fibroblasts 2 Markers
Macrophages 1 Markers
Macrophages 2 Markers
Macrophages 3 Markers
Macrophages 4 Markers
Neurons Markers
Plasmacytoid DCs Markers
VSMCs 1 Markers
VSMCs 2 Markers
VSMCs 3 Markers
VSMCs 4 Markers
VSMCs 5 Markers
VSMCs 6 Markers
VSMCs 7 Markers
wnnUMAP_1
wnnUMAP_2
1.0
0.5
0.0

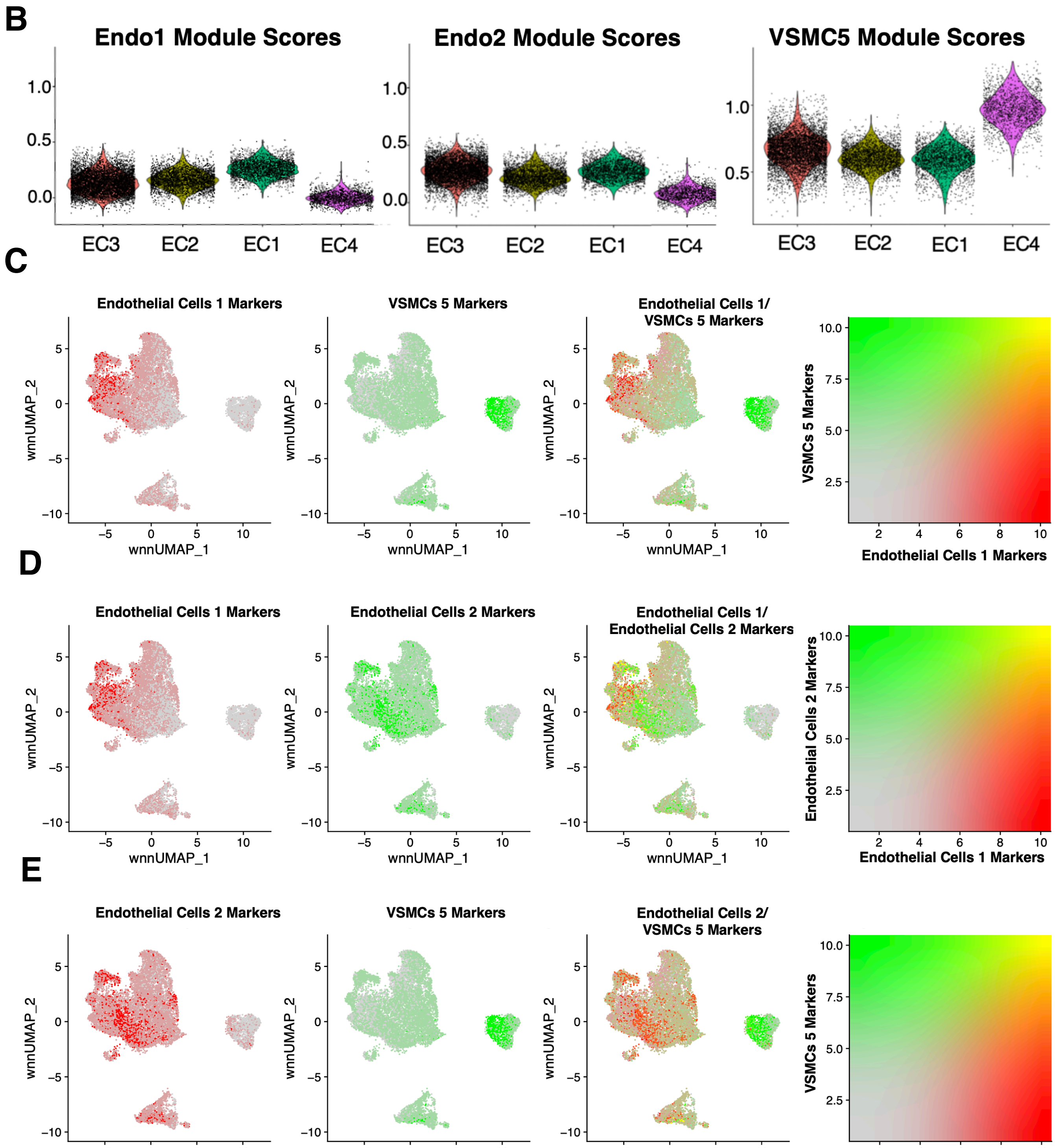


**Supplementary Figure 7 (Figure S7) – Related to Figure 5 | (A)**, Feature plots for each *ex vivo* module score across *in vitro* cells. Briefly, *ex vivo* module scores are generated using top marker genes for each *ex vivo* cell subtype. The Seurat function AddModuleScore is used to score each cell for visualization. **(B)**, Violin plots displaying Endothelial Cells 1 (Endo1), Endothelial Cells 2 (Endo2), and VSMCs 5 (VSMC5) module scores for each perturbation across *in vitro* EC1-4. **(C)**, Feature plots displaying distribution of Endo1 (red) versus Endo2 (green) module scores across *in vitro* cells. **(D)**, Feature plots displaying distribution of Endo1 (red) versus VSMC5 (green) module scores across *in vitro* cells. **(E)**, Feature plots displaying distribution of Endo2 (red) versus VSMC5 (green) modules scores across *in vitro* cells.

A

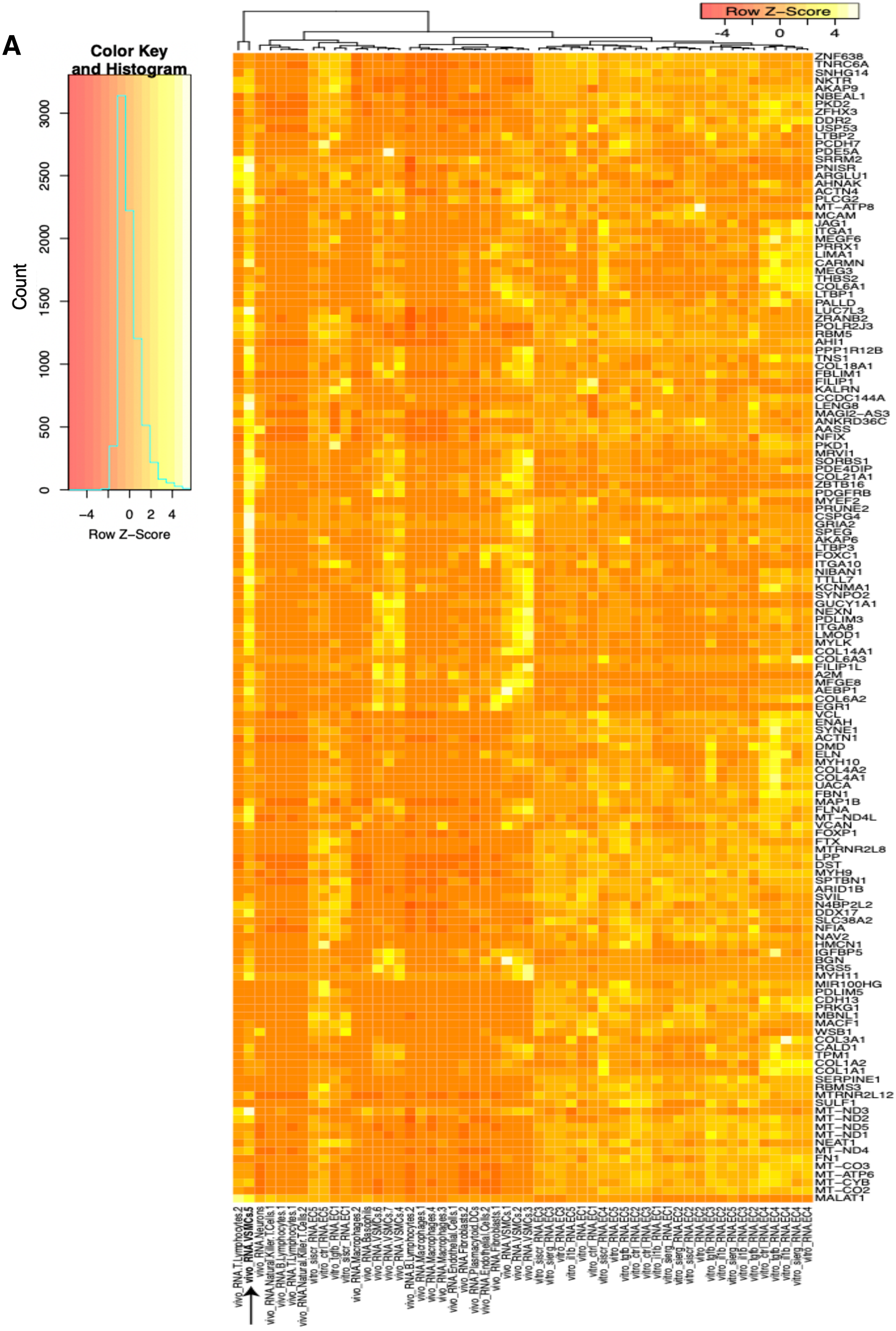
Color Key and Histogram
Count
Row Z-Score
Row Z-Score
ZNF638
TNRC6A
SNHG14
NKTR
AKAP9
NBEAL1
PKD2
ZFHX3
DDR2
USP53
LTBP2
PCDH7
PDE5A
SRRM2
PNISR
ARGLU1
AHNAK
ACTN4
PLCG2
MT-ATP8
MCAM
JAG1
ITGA1
MEGF6
PRRX1
LIMA1
CARMN
MEG3
THBS2
COL6A1
LTBP1
PALLD
LUC7L3
ZRANB2
POLR2J3
RBM5
AHI1
PPP1R12B
TNS1
COL18A1
FBLIM1
FILIP1
KALRN
CCDC144A
LENG8
MAGI2-AS3
ANKRD36C
AASS
NFIX
PKD1
MRVI1
SORBS1
PDE4DIP
COL21A1
ZBTB16
PDGFRB
MYEF2
PRUNE2
CSPG4
GRIA2
SPEG
AKAP6
LTBP3
FOXC1
ITGA10
NIBAN1
TTLL7
KCNMA1
SYNPO2
GUCY1A1
NEXN
PDLIM3
ITGA8
LMOD1
MYLK
COL14A1
COL6A3
FILIP1L
A2M
MFGE8
AEBP1
COL6A2
EGR1
VCL
ENAH
SYNE1
ACTN1
DMD
ELN
MYH10
COL4A2
COL4A1
UACA
FBN1
MAP1B
FLNA
MT-ND4L
VCAN
FOXP1
FTX
MTRNR2L8
LPP
DST
MYH9
SPTBN1
ARID1B
SVIL
N4BP2L2
DDX17
SLC38A2
NFIA
NAV2
HMCN1
IGFBP5
BGN
RGS5
MYH11
MIR100HG
PDLIM5
CDH13
PRKG1
MBNL1
MACF1
WSB1
COL3A1
CALD1
TPM1
COL1A2
COL1A1
SERPINE1
RBMS3
MTRNR2L12
SULF1
MT-ND3
MT-ND2
MT-ND5
MT-ND1
NEAT1
MT-ND4
FN1
MT-CO3
MT-ATP6
MT-CYB
MT-CO2
MALAT1
vivo_RNA.T.Lymphocytes.2
vivo_RNA.VSMCs.5
vivo_RNA.Neurons
vivo_RNA.Natural.Killer.T.Cells.1
vivo_RNA.B.Lymphocytes.1
vivo_RNA.T.Lymphocytes.1
vivo_RNA.Natural.Killer.T.Cells.2
vitro_siscr_RNA.EC5
vitro_ctrl_RNA.EC5
vitro_tgfb_RNA.EC1
vitro_siscr_RNA.EC1
vivo_RNA.Macrophages.2
vivo_RNA.Basophils
vivo_RNA.VSMCs.6
vivo_RNA.VSMCs.7
vivo_RNA.VSMCs.4
vivo_RNA.B.Lymphocytes.2
vivo_RNA.Macrophages.1
vivo_RNA.Macrophages.4
vivo_RNA.Macrophages.3
vivo_RNA.Endothelial.Cells.1
vivo_RNA.Fibroblasts.2
vivo_RNA.Plasmacytoid.DCs
vivo_RNA.Endothelial.Cells.2
vivo_RNA.Fibroblasts.1
vivo_RNA.VSMCs.1
vivo_RNA.VSMCs.2
vivo_RNA.VSMCs.3
vitro_siscr_RNA.EC3
vitro_sierg_RNA.EC3
vitro_RNA.EC3
vitro_il1b_RNA.EC5
vitro_RNA.EC1
vitro_ctrl_RNA.EC1
vitro_siscr_RNA.EC4
vitro_RNA.EC5
vitro_tgfb_RNA.EC5
vitro_ctrl_RNA.EC2
vitro_ctrl_RNA.EC3
vitro_il1b_RNA.EC1
vitro_sierg_RNA.EC1
vitro_sierg_RNA.EC2
vitro_siscr_RNA.EC2
vitro_RNA.EC2
vitro_tgfb_RNA.EC3
vitro_il1b_RNA.EC2
vitro_sierg_RNA.EC5
vitro_il1b_RNA.EC3
vitro_tgfb_RNA.EC2
vitro_ctrl_RNA.EC4
vitro_tgfb_RNA.EC4
vitro_il1b_RNA.EC4
vitro_sierg_RNA.EC4
vitro_RNA.EC4

**B**

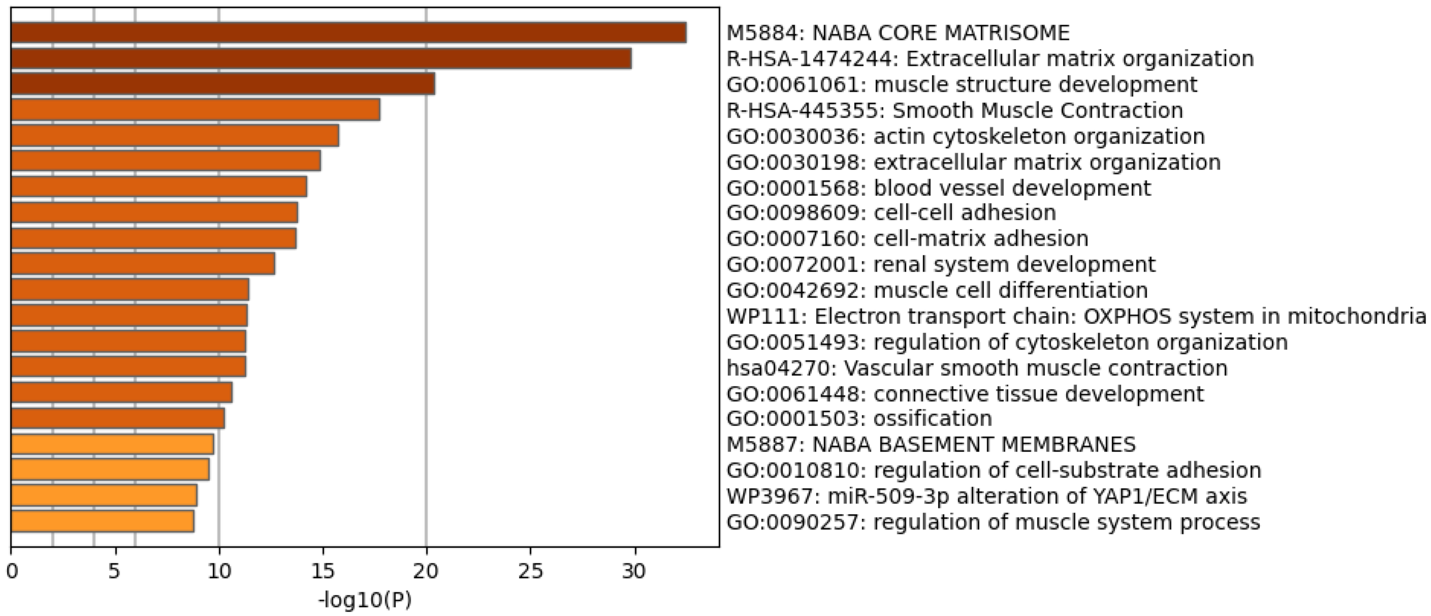


**Supplementary Figure 8 (Figure S8) – Related to Figure 5 I (A)**, Heatmap displaying average expression of VSMC5 maker genes (black arrow) across *in vitro* and *ex vivo* datasets. Rows (genes) and columns (cell subtypes) are clustered based on average expression for each given gene. **(B)**, PEA of the top 200 genes for VSMC5 (adjusted p-value < 0.05).

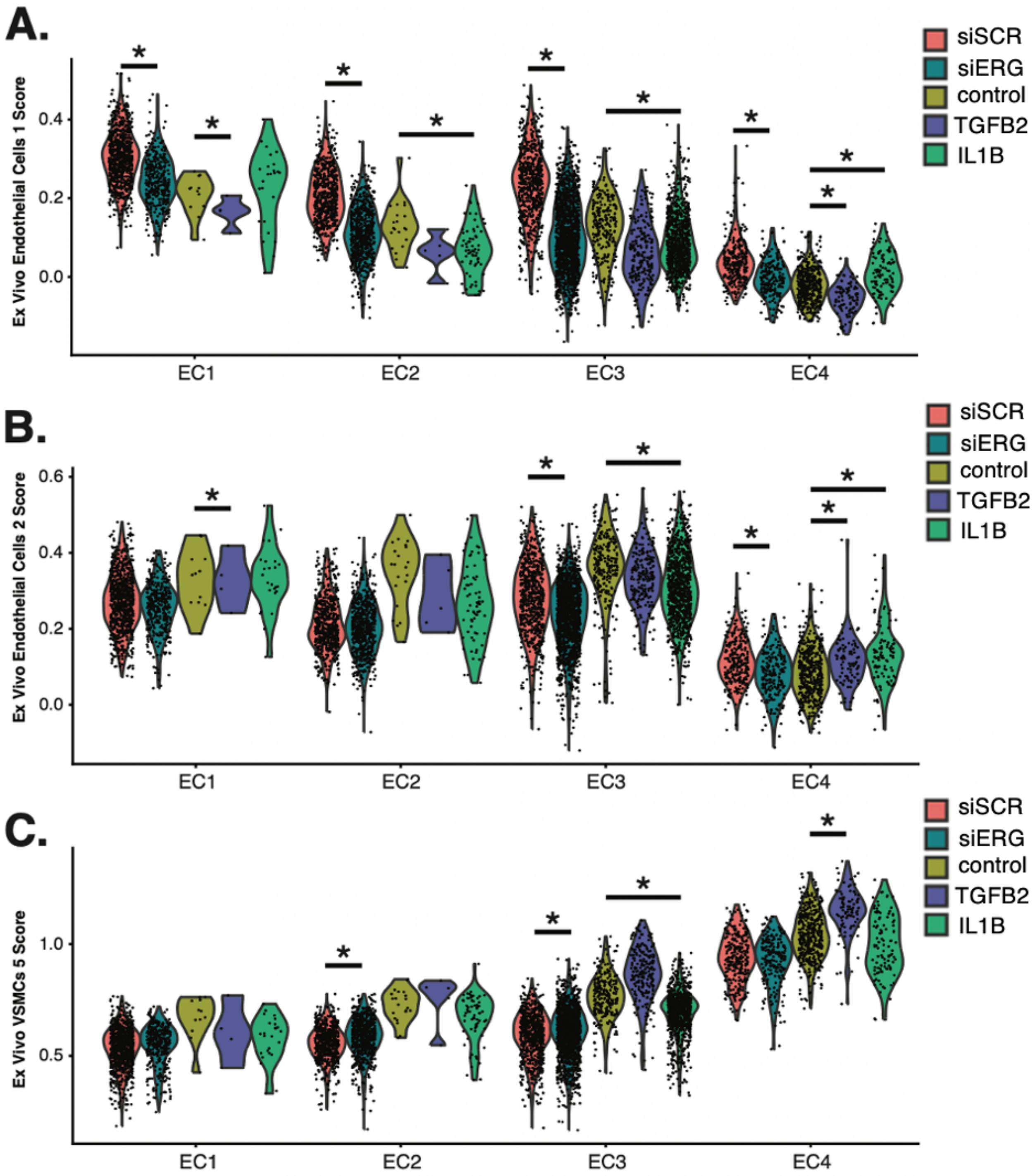


**Supplementary Figure 9 (Figure S9) – Related to Figure 5 I (A)**, Violin plots of *ex vivo* Endo1 module scores across EC1-4. **(B)**, Violin plots of *ex vivo* Endo2 module scores across EC1-4. **(C)**, Violin plots of *ex vivo* VSMC5 module scores across EC1-4. Adjusted p-value for A-C generated using Wilcoxon rank sum test with continuity correction by setting the alternative hypothesis to "two.sided".

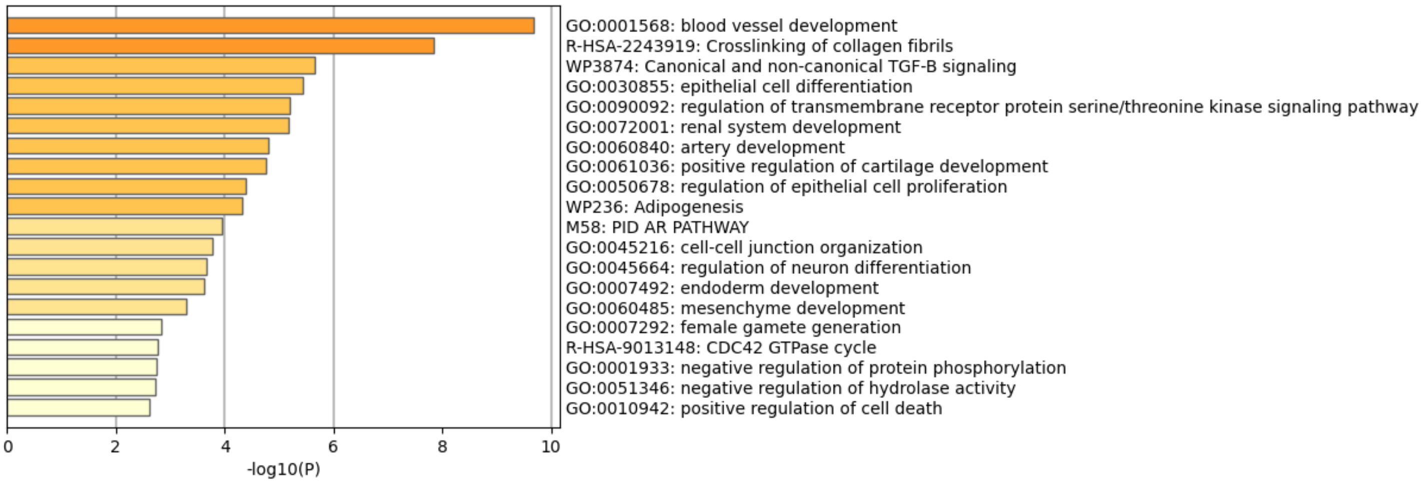


**Supplementary Figure 10 (Figure S10) – Related to Figure 6 l** PEA of significant (p-value < 0.05) EC4 linked genes which overlap with significant (p-value < $10^{-8}$) CAD associated SNPs.

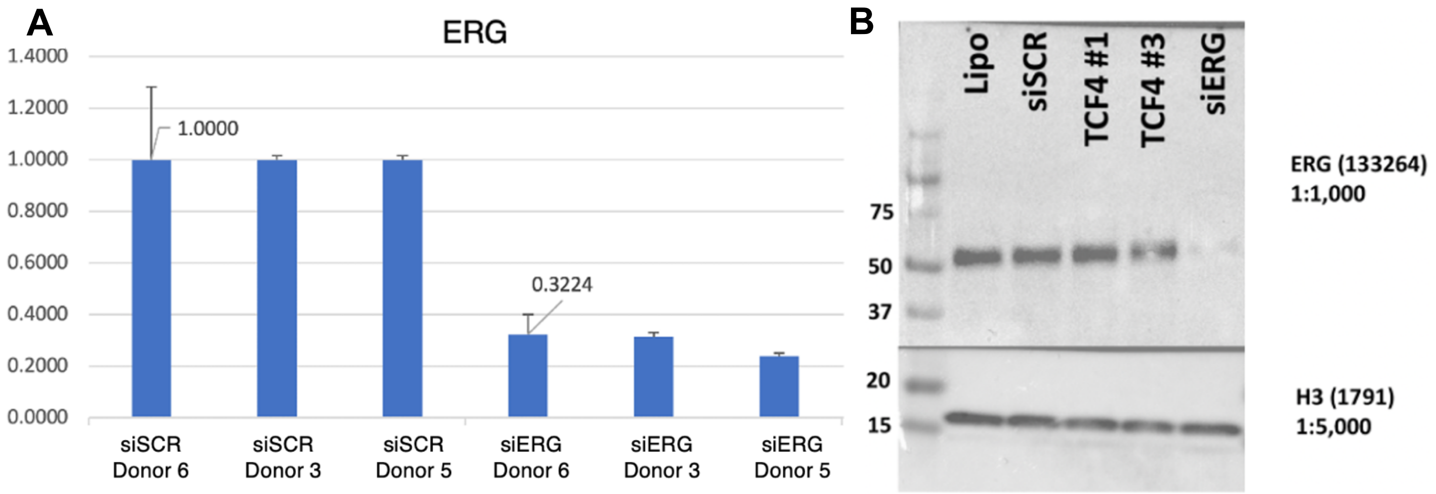


**Supplementary Figure 11 (Figure S11) – Related to Methods l (A)**, qPCR results for ERG knockdown across donors. **(B)**, Western Blot representing a typical knockdown of siERG with the siRNA pools used in this study. TCF4 samples are irrelevant for the purposes of this study.

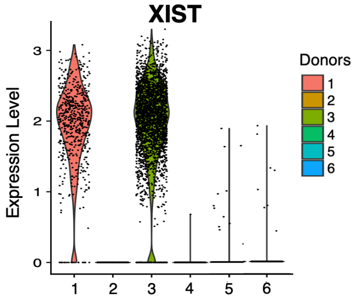


**Supplementary Figure 12 (Figure S12) – Related to Methods l** Violin plot of *XIST* showing expected expression in female *in vitro* donor cells (1 and 3) and lack of expression in male *in vitro* donor cells (2, 4, 5, and 6).
